# Synchronous 3D patterning of diverse CNS progenitors generates motor neurons of broad axial identity

**DOI:** 10.1101/2024.12.13.628384

**Authors:** Felix Buchner, Zeynep Dokuzluoglu, Joshua Thomas, Antonio Caldarelli, Shrutika Kavali, Fabian Rost, Tobias Grass, Natalia Rodríguez-Muela

## Abstract

In vitro human organoid models have become transformative tools for studying organogenesis, enabling the generation of spinal cord organoids (SCOs) that mimic aspects of spinal cord biology. However, current models do not produce spinal motor neurons (spMNs) with a wide range of axial identities along spinal cord segments within a single structure, limiting their utility in understanding human neural axial specification and the selective vulnerability of spMN subpopulations in motor neuron diseases. Here, we present a novel approach to enhance spMN axial heterogeneity in an advanced SCO model derived from neural stem cells (NSCs) and retinoic acid (RA)-primed neuromesodermal progenitors (NMPs). RA priming guided NMP differentiation into caudal neural progenitors, generating SCOs enriched in spMNs with posterior axial identities. To further diversify spMN populations, we optimized differentiation by synchronously patterning NSCs with RA-primed NMPs. Incorporating an endothelial-like network and skeletal muscle cells enhanced the organoids’ physiological complexity and neural maturation and organoid cell viability. This comprehensive approach, termed CASCO, provides a robust platform to study human spMN specification and model neurodegenerative diseases.

## Introduction

Advancements in 3D in vitro modelling of the human central nervous system hold great promise to help understand human physiology and pathology of the brain and the spinal cord. Following established protocols on human induced pluripotent stem cells (hiPSCs), researchers have successfully constructed region-specific organoids, including those of the retina, hippocampus, thalamus, midbrain, and cerebellum (reviewed in Guy et al. 2021, Mulder et al. 2023). Recent progress has also enabled the directed differentiation of stem cells into spinal cord-like organoids (SCOs). However, replicating the molecular diversity of spinal motor neurons (spMNs) found in vivo in a single organoid has not been achieved. spMNs are cholinergic neurons located in the ventral horn of the spinal cord and are responsible for muscle contraction and motor function. Developing such an organoid would be instrumental to dissect the plethora of events orchestrating axial spMN specification during human spinal cord development and the underlying mechanisms of the selective degeneration of specific spMN subpopulations occurring in motor neuron diseases. Several approaches have successfully patterned pluripotent stem cells (PSCs) into 2D neural spinal progenitors and spinal spheroids using chemically defined media containing specific patterning molecules that mimic key spinal cord developmental steps. The first studies used dual SMAD inhibition to block Activin/nodal and BMP signalling pathways of the TGFß superfamily, which generated a high percentage of PAX6^+^ NSCs (Chambers et al. 2009, Surmacz et al. 2012). These cells were then exposed to retinoic acid (RA) – which induces rostral Homeobox (HOX) gene expression (Kawanishi et al. 2003, Joksimovic et al. 2005, Mazzoni et al. 2013) – and Sonic hedgehog agonists – which direct neural progenitors toward a ventral spinal cord fate (Wicherle et al. 2002, Stanton et Peng 2010, Dessaud et al. 2010, Ogura et al. 2018) – to imitate developmental signals released from somites (RA) and Floor Plate/notochord (SHH) during neural tube development in vivo (Wichterle et al. 2002, Meinhardt et al. 2014, Davis-Dusenbery 2014). While we and others have applied similar guided differentiation approaches to in vitro modelling of diseases affecting spMNs (Rodriguez Muela et al. 2017, Rodriguez Muela et al. 2018, Hor et al 2018, Baxi et al. 2022), the existing protocols are limited at generating a wide heterogeneity of spMNs along the rostro-caudal (RC) axis (Buchner et al. 2023).

Neuromesodermal progenitors (NMPs) are a bipotent cell population located in the caudal lateral epiblast zone of the developing embryo. They are crucial for the axial extension of the embryo and secondary neurulation, as they contribute to both neuroectoderm and mesoderm lineages (Shaker et al. 2021). NMPs are developmentally primed to generate CNS caudal structures, which include the spinal cord and musculoskeletal system (Gouti et al. 2014; Tsakiridis et al. 2014, Faustino-Martins et al. 2020). It has been recently demonstrated that the establishment of an axial identity occurs before further lineage allocation during spinal cord development (Metzis et al. 2018) and that key signaling pathways regulate the molecular identity, emergence and function of NMPs, balancing their self-renewal and differentiation capacity (Gouti et al. 2014). Gradients of WNT, FGFs and GDF11 regulate *HOX* gene expression during body axis elongation during early embryo development (Ciruna et Rossant 2001; Forlani et al. 2003, Mouilleau et al. 2021). Most importantly, these pathways induce the expression of the transcription factor *CDX2* (Nordström et al. 2006), an upstream regulator of central and posterior *HOX* genes (Keenan et al. 2006; Gaunt et al. 2004; Gaunt et al. 2017; Neijts et al. 2017) key for axial elongation (Amin et al. 2016) that is suppressed when stem cells have committed to an anterior neural fate (Nordström et al. 2006, Metzis et al. 2018). Similarly, in vivo experiments have demonstrated that GDF11, accumulating in the tailbud region, is indispensable for the correct anatomical formation of the different axial segments of the spinal cord and skeleton (Jurberg et al. 2014), and specifically instructs *HOX8-HOX10* gene expression in the thoracic and lumbar spinal cord, as well as *HOX13* expression in the most caudal region of the tailbud (Liu et al. 2001; Liu 2006; Andersson et al. 2006, Aires et al. 2019). Furthermore, in vivo, RA has been shown to guide NMPs toward an NSC fate (Gouti et al. 2014; Shaker et al. 2021). By modulating the concentration and timing of RA exposure to NMPs in vitro, researchers can guide the regional identity of the developing neurons toward caudal fates, which is crucial for modeling lower spinal cord regions, corresponding to the vertebral levels L1 to S5 (Lippmann et al. 2015; Mouilleau et al. 2021). The RC axial identity of spMNs is determined by the *HOX* gene expression code (Soshnikova et al. 2013). *HOX* genes are expressed in a collinear fashion along the RC axis, with their genomic order corresponding to their spatial expression in the developing embryo. This arrangement is critical for establishing the positional identity of spMNs, ensuring that they acquire characteristics appropriate to their location along the spine (Deschamps et al. 2017). On a molecular level, RA also plays a key role in this process by firstly promoting primary neurogenesis through inducing the expression of proneural genes such as *NEUROD1* (Maden et al. 1996, Diez del Corral 2003, Janesick et al. 2015, Haushalter et al. 2017) and, secondly, by activating anterior *HOX* genes in the cervical region of the neural tube, targeting specific genomic sites within the *HOX4-5* gene cluster (Mahony et al. 2011).

Several SCO models have used NMPs to produce a variety of cell types that mimic the early stages of spinal cord development, serving as a foundational component to replicate the complex architecture of the human spinal cord (reviewed in Buchner et al. 2023, Zhou et al. 2024). However, recent in vitro patterning protocols aimed at generating SCOs starting from NMPs often encounter significant challenges. Due to the low amount of RA used in these protocols, these approaches frequently result in a limited number of spMNs, while retaining a high proportion of progenitor cells. Additionally, the use of NMPs as a starting cell population frequently leads to the production of non-neuronal lineages, such as muscle cells, due to the bipotent nature of these cells. According to the knowledge gained through developmental neurobiology studies on model organisms, controlling axial allocation fates should be possible by adjusting the culture time of NMPs in media containing WNT agonist, FGF2/8b and GDF11 before RA stimulation (Lippmann et al. 2015). Early exposure of NPCs or NMPs to this growth factor cocktail advances HOX clock progression towards caudal spMN fate allowing the cells to maintain their established axial identity even after neural differentiation (Mouilleau et al. 2021; Metzis et al. 2018, Grass et al. 2024). A recent study reported the in vitro generation of spMNs across the entire RC axis within embryo-like structures using FISH against *HOX* transcripts as readout; however, this work primarily focused on the morphogenetic processes during the elongation of the model, and the specific anatomical identity of the generated spMNs was not characterized (Gribaudo et al. 2023).

Most organoid platforms to model neuromuscular diseases have a neuroscentric focus and lack the cell lineage diversity found in vivo, such as skeletal muscle cells and a functional vasculature. Without vasculature, organoids struggle with inefficient nutrient delivery and waste removal, leading to necrotic cores and limited size expansion (Cakir et al. 2019, Buchner et al. 2023). The contribution of vasculature-derived signalling molecules to developmental neurogenesis, as well as to neuronal maturation and the establishment of functional neural networks is also proven (Vogenstahl et al. 2022, Wimmer et al. 2019, Ng et al. 2021, Cakir et al. 2019). Similarly, without skeletal muscle cells, spMNs in the organoids lack their physiological target, as well as the mechanical forces and signalling cues provided by the muscle, which are essential for proper tissue function (Faustino-Martins et al. 2021). Incorporating a functional endothelial network and skeletal muscle elements should thus enhance nutrient exchange, support sustained growth and provide the necessary biomechanical environment, thus ultimately improving the physiological relevance of the model for studying human physiology and disease (Matthews et al. 2009, Iyer et al. 2022, Sun et al. 2022).

Building on the existing literature on mouse and human spinal cord development, as well as on studies on in vitro spMN differentiation from PSCs, we developed an advanced protocol to generate SCOs composed by spMNs with a balanced RC diversity embedded in a physiologically relevant environment. This new method starts by shortly exposing hiPSC-induced NMPs to RA to prime them to efficiently commit towards the neural lineage. These primed NMPs exhibited a strong bias for neuroectoderm over mesodermal lineage formation and showed high efficiency in differentiating into caudal neural progenitors. Using these neuroectoderm-poised NMPs as starting stem cell source, we designed and screened 12 differentiation protocols to identify the most effective method at generating a diverse array of RC spMN subpopulations. We further optimized this method by subjecting an even mix of NSCs - which, under our culture conditions, commit to anterior regions of the spinal cord- and caudally-primed NMPs as starting stem cell pool to the selected differentiation protocol. To the best of our knowledge, this approach produces SCOs composed of a high abundance of spMNs with the widest axial heterogeneity reported to date. The controlled and synchronous development of functional vasculature-like structures and skeletal muscle cells intertwined with the neural tissue added physiological relevance of this model. Our novel comprehensive axial spinal cord organoid (CASCO) approach for SCO formation sets the stage for more accurate human in vitro modelling of spinal cord development and provides a platform to investigate the selective degeneration of specific spMN subtypes in neuromuscular diseases.

## Results

### Commonly applied paradigms for hiPSC-derived spMN formation mostly produce neurons with rostral identity

Most existing protocols for generating spMNs from hiPSCs are adaptations of the original method established by Wichterle et al. (Cell, 2002) using mouse embryonic stem cells. This field-standard protocol, which induces the formation of NSCs through dual SMAD inhibition via SB431542 (SB) and LDN193189 (LDN), and relies on an early and extended exposure of high RA concentration to promote ventralization, results predominantly in the formation of rostral spMNs (Wichterle et al. 2002; Surmacz et al. 2012; Amoroso et al. 2013) (Fig. 1A). Given our objective to develop a procedure capable of specifying spMNs across the entire RC axis, we used this method as a "rostral" benchmark for comparison against our new protocols. Following this protocol, we patterned a healthy hiPSC line (BJ-siPS) into spinal spheroids, which abundantly expressed the spMN progenitor marker OLIG2 and the spMN marker ISL1/2 by day 20 of differentiation (Fig. 1B). As expected, single-cell RNA sequencing showed that these spheres are predominantly composed of rostral spMNs. Six main cell clusters were identified (Fig. 1C-D). The majority of cells expressed pan neuronal markers such as *MAP2, STMN1/2* and *ELAV4* and early spMN development markers, such as *ONECUT2*; only ∼5% remained progenitors in a proliferative state, expressing the cell cycle markers *TOP2A*, *MKI67* or *CDK1* (Fig. 1C-F). A subset of cells (10-20%) expressed canonical spMN identity markers, *MNX1(HB9), ISL1* or *CHAT*. 25% of all cells showed a transcriptomic profile compatible with cervical/cranial spMN identity, expressing *MEIS1, ALCAM* and *PHOX2A/B* (Sagner et al 2018, Sagner et al. 2021) (Fig. 1D-F), and more than 90% of cells expressed rostral *HOX* genes like *HOXB4* or *HOXA5* (Fig. 1D-E, G). Only 10% of cells adopted a more caudal, brachial-thoracic identity, expressing *HOXC6*, *HOXC8* or *HOXC9*. No cells expressed lumbar *HOXA10* or *HOXC10* genes (Fig. 1D). In summary, this NSC-based protocol is efficient at generating rostral spMNs, but does not enable the formation of spMNs corresponding to lumbar spinal cord segments, which originate from NMPs. Therefore, for our advanced protocol we decided to optimize the initial steps of stem cell patterning and used NMPs as a starting stem cell pool, with the goal of producing a more varied axial spMN diversity.

**Figure 1.**
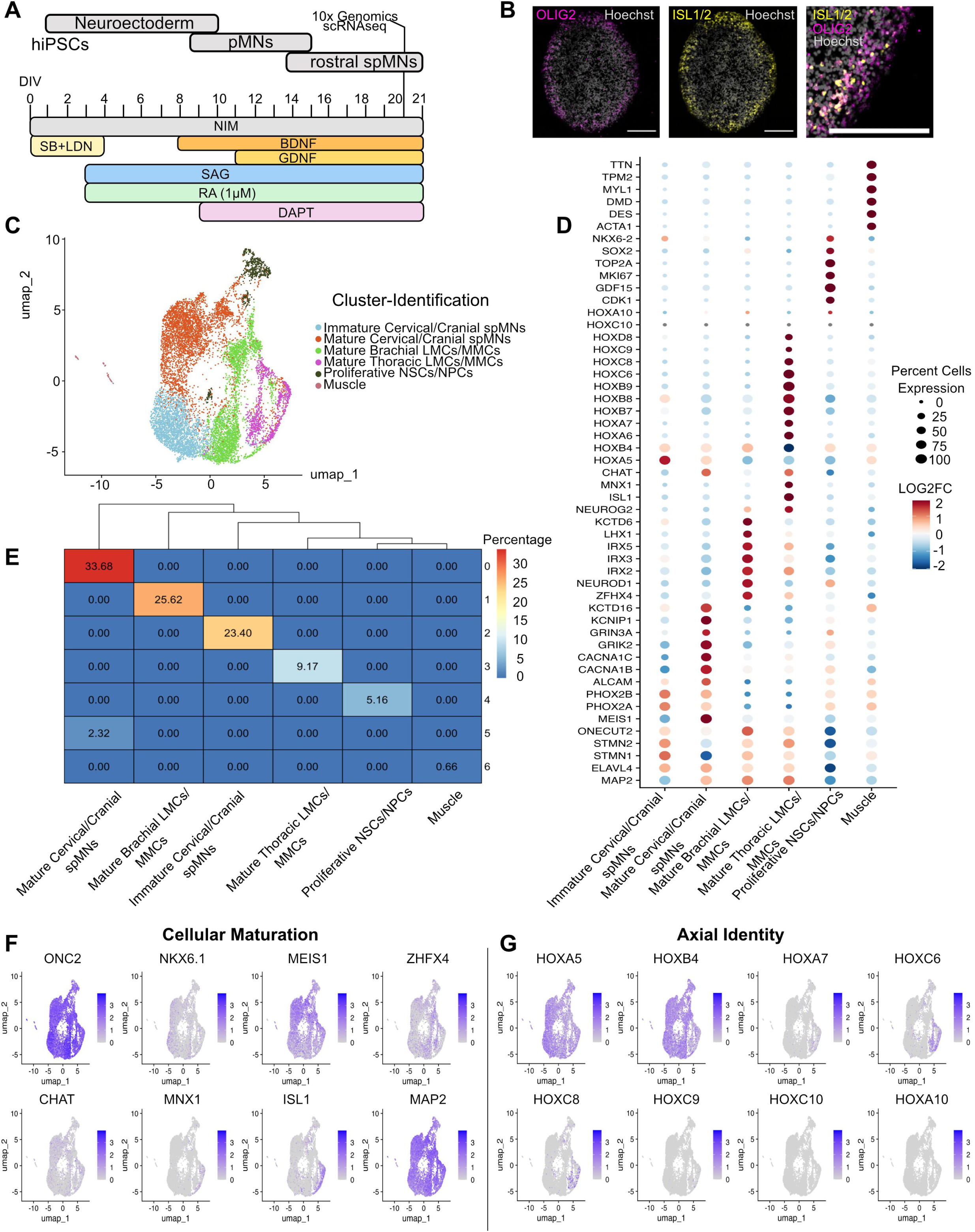
Commonly applied paradigms for hiPSC-derived spMN formation mostly produce spMNs with rostral identity. **A.** Scheme showing the commonly used protocol to differentiate hiPSCs into ventral spMNs through the generation of NSCs. **B.** Immunostained hiPSC-NSC derived 20 day spheres showing immunoreactivity against pMN marker OLIG2 and spMN marker ISL1/2. Scale bar is 200 µM. **C.** UMAPs projections of 12.637 sequenced cells isolated from hiPSC-NSC derived 20 day spheres, subjected to 10xGenomics-based scRNAseq.. **D.** Dotplot showing LOF2FC of marker gene expression for each cellular cluster that was previously identified in the scRNAseq analysis. **E.** Confusion matrix of all sequenced cells, showing relative percentage of each cell cluster. **F.** UMAPs showing early maturation stage of spMNs and neurons within the spinal spheres. **G.** UMAPs showing axial identity of hiPSC-NSC derived 20 day spheres

### hiPSC-derived NMPs primed by retinoic acid efficiently generate caudal neuroectoderm

We and others have shown that hiPSC-derived NMPs constitute a powerful source to generate neuromuscular organoids (Faustino-Martins et al. 2020, Pereira et al. 2021, Grass et al. 2024, Gao et al. 2024). As previously introduced, the molecular identity of NMPs is safeguarded by overlapping morphogen activity, including WNT, FGF and RA signalling. Importantly, loss of RA activity disrupts the cell fate balance in the NMP niche, leading to increased *TBX6*^+^ mesoderm near the primitive streak, reduced *SOX2* expression in the caudal epiblast and overall impaired neural differentiation (Cunningham et al. 2011, Cunningham et al. 2015, Javali et al. 2017). To generate NMPs poised to commit to a neural lineage, we exposed hiPSCs to WNT agonist CHIRON99021 (CHIR) and FGF2 for 3 days according to previous studies (Gouti et al. 2014, Cunningham et al. 2015), then exposed them for 2 hours to RA (Horton et Maden 1995) and analyzed the samples the next day. We compared these spheres with a separate group generated following the standard NSC-based protocol described above (SB+LDN) (Fig. 2A). This new approach resulted in the efficient generation of spheres composed of SOX2^Low+^:CDX2^+^ and PAX6^Low+^:TBXT^+^ NMPs, while the spheres generated from the NSC-based protocol were mostly comprised of SOX2^+^ and PAX6^+^ cells (Fig. 2B-C). Although SOX2 transcript levels were only slightly decreased in the NMP condition relative to SB+LDN cultures, we observed a clear decrease in the number of SOX2^+^ cells. NMP-condition cultures also exhibited an increase in the levels of NMP-specific transcripts TBXT, GBX2, NT5E and WNT3A (Fig. 2D), and acquired a different cell morphology (Fig. 2E).

**Figure 2.**
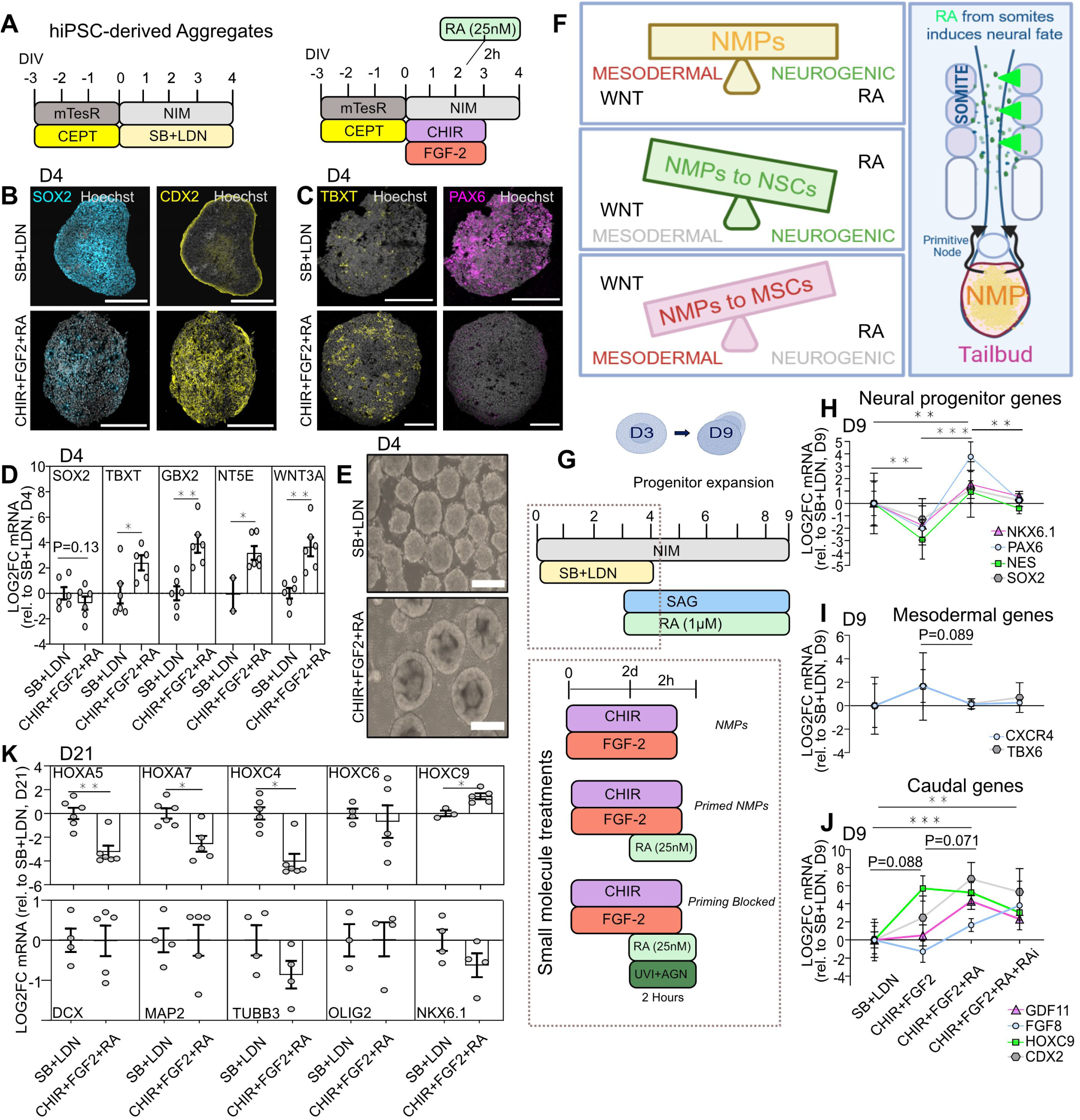
hiPSC-derived NMPs primed by retinoic acid efficiently generate caudal neuroectoderm. **A.** Scheme showing the approach by which hiPSCs were pre-patterned into NMPs. **B.** Immunostaining of day 4 SB+LDN or CHIR+FGF-2+RA generated aggregates against the NSC marker SOX2 and caudal epiblast marker CDX2. **C.** Immunostaining of day 4 SB+LDN or CHIR+FGF-2+RA generated aggregates against the mesodermal TBXT and the proneural PAX6. **D.** qRT-PCR on day 4 spinal spheres generated according to the 4 paradigms shown in (G) showing NMP markers *SOX2*, *TBXT*, *GBX2*, *NT5E*, and *WNT3A* (Unparametric Mann-Whitney-U test, N=2 differentiations with n=1-3 technical replicates, each with 5-10 pooled spheres. Each dot is a technical replicate). **E.** Representative bright field images of NSC (SB+LDN) and axial progenitor (CHIR+FGF-2+RA) aggregates at day 4. Scale bar, 500 µM. **F.** Scheme showing NMPs’ bipotency to develop into either neural cells or mesodermal lineages, which is mediated through modulation of WNT and RA pathway activation. **G.** Scheme showing timeline of different patterning approaches followed to generate spinal spheres. **H.** qRT-PCR on day 9 spinal spheres generated according to the 4 paradigms shown in (G) showing expression of NSC genes (*PAX6, NES, NKX6.1, SOX2*). (One-way ANOVA with Fishers LSD or unparametric Mann-Whitney-U test N=4 differentiations with n=3 technical replicates, each with 5-10 pooled spheres). **I.** Gene expression of mesodermal markers (*CXCR4, TBX6*) from similar spheres as in H. **J.** Gene expression of caudal genes (*FGF8, HOXC9, GDF11, CDX2*) from similar spheres as in H. **K.** qRT-PCR on day 21 spinal spheres generated according to the 4 paradigms shown in (G) showing caudal HOXs (HOXC6, HOXC9), rostral HOX transcripts (*HOXA5, HOXA7, HOXC4*), and neuronal identity makers (*DCX, MAP2, TUBB3, OLIG2, NKX6.1*). (Unparametric Mann-Whitney-U test, N=2 differentiations with n=1-3 technical replicates, each with 5-10 pooled spheres). For all graphics: Mean+SEM are shown, *p < 0.05, **p < 0.01, ***p < 0.001.

The primary challenge when generating SCOs from NMPs as a starting population is to refine the differentiation protocols to effectively steer these progenitor cells toward neural lineage commitment by improving neural progenitor expansion, while minimizing their tendency to differentiate into mesodermal lineages (Fig. 2F). This can be achieved by tuning their exposure to WNT signalling and RA (Cunningham et al. 2015, Patani et al. 2011). To evaluate the efficacy of RA in guiding the differentiation trajectory of NMPs towards ventral spinal cord neural lineage, we cultured the spheres for 6 additional days and exposed them to different paradigms: NSC induction (SB+LDN), NMP induction (CHIR+FGF2), NMP induction with RA exposure (CHIR+FGF2+RA) and as a treatment control, NMP induction with RA and RA inhibitor treatment (UVI/AGN) (Fig. 2G). qPCR analysis from the differently cultured spheres revealed a reduced expression of neural progenitor markers (*SOX2*, *PAX6*, *NES*, *NKX6.1*) in the NMP condition (CHIR+FGF2) compared to the NSC spheres (SB+LDN), and highest expression levels in the RA-primed NMPs (CHIR+FGF2+RA) (Fig. 2H). This effect was specific to RA pulse, as evidenced by its reversal with RA inhibitors (CHIR+FGF2+RA+RAi). The NMP condition still showed an increase, yet non-significant, in mesodermal transcripts (*CXCR4*, *TBX6*) that was abolished in the RA primed-NMP condition (Fig. 2I). As expected, the NMP spheres and RA-primed NMP spheres expressed more caudal transcripts (*FGF8*, *HOXC9*, *GDF11*, *CDX2*) than the NSC spheres (Fig. 2J) and, importantly, exposure to RA or RA+RAi did not significantly alter their caudal identity. Moreover, interestingly, *FGF8* levels appeared higher in the CHIR+FGF2+RA+RAi condition compared to RA alone, matching in vivo studies which showed that RA counteracts *FGF8* expression (Kumar et Duester 2014). Next, we examined the different spheres at day 21 to explore whether the initial neural-bias introduced by the RA pulse on the NMPs was maintained as the progenitors developed and whether this was sufficient to generate caudal neurons. qPCR analysis of *HOX* genes on day 21 spheres revealed a shift from rostral (*HOXA5, HOXA7, HOXC4, HOXC6*) to caudal transcripts (*HOXC9*) in the RA-primed NMP condition compared to the NSC condition (Fig. 2K). No significant difference was observed in any of the neural genes measured (*DCX*, *MAP2*, *TUBB3*, *OLIG2*, *NKX6.1*) (Fig. 2K). Together, these results indicate that priming NMPs with a brief RA pulse to direct their neural lineage commitment is sufficient to generate neurons at levels comparable to NSCs, without interfering with the caudal features of NMPs, and significantly enhances caudal HOX gene expression.

### Neuro-committed RA-primed NMPs generate thoracic and lumbar spMNs independently of the patterning approach followed

During early development, the mammalian embryo undergoes axial extension, with axial progenitors continuously proliferating and undergoing neural induction due to RA diffusing from the somites (Deschamps et al. 2017, Shaker et al. 2021). The timing of neural induction and caudalization of the tail bud region is closely interconnected, as the extension of the body axis through NMP proliferation during embryogenesis coincides with the secretion of neuronal fate-inducing molecules from the surrounding tissues, such as the notochord and presomitic mesoderm (Amin et al. 2016). However, it is unclear when NMPs need to be exposed to proliferative, neural inductive and caudalizing signals in vitro to expand their potential to efficiently generate caudal spMNs. To shed light on this problem, we designed an approach to systematically test 11 different differentiation protocols, divided into three modules (Fig. 3A, S1A and Supplemental table 1): Module 1, simultaneous neuroprogenitor expansion/caudalization and neuroectoderm induction; Module 2, neuroectoderm induction followed by neuroprogenitor expansion/caudalization and Module 3, neuroprogenitor expansion/caudalization prior to neuroectoderm induction. A key difference between these protocols is when the neuroectoderm induction phase using SB+LDN (Chambers et al. 2009) is applied, either during, before or after caudalization signals, respectively. Patterning protocol #1 was used as control in which RA-primed NMPs were subjected to the field-standard spMN differentiation protocol (Fig. 1A), while patterning protocols #2 to #12 were tested to develop an improved method to effectively generate spMNs with a more diverse RC identity (Fig. S1A). For this purpose, we conducted a comparative assessment of neural, spMN and *HOX* expression levels at multiple time points corresponding to different stages of SCO development (Fig. 3B).

**Figure 3.**
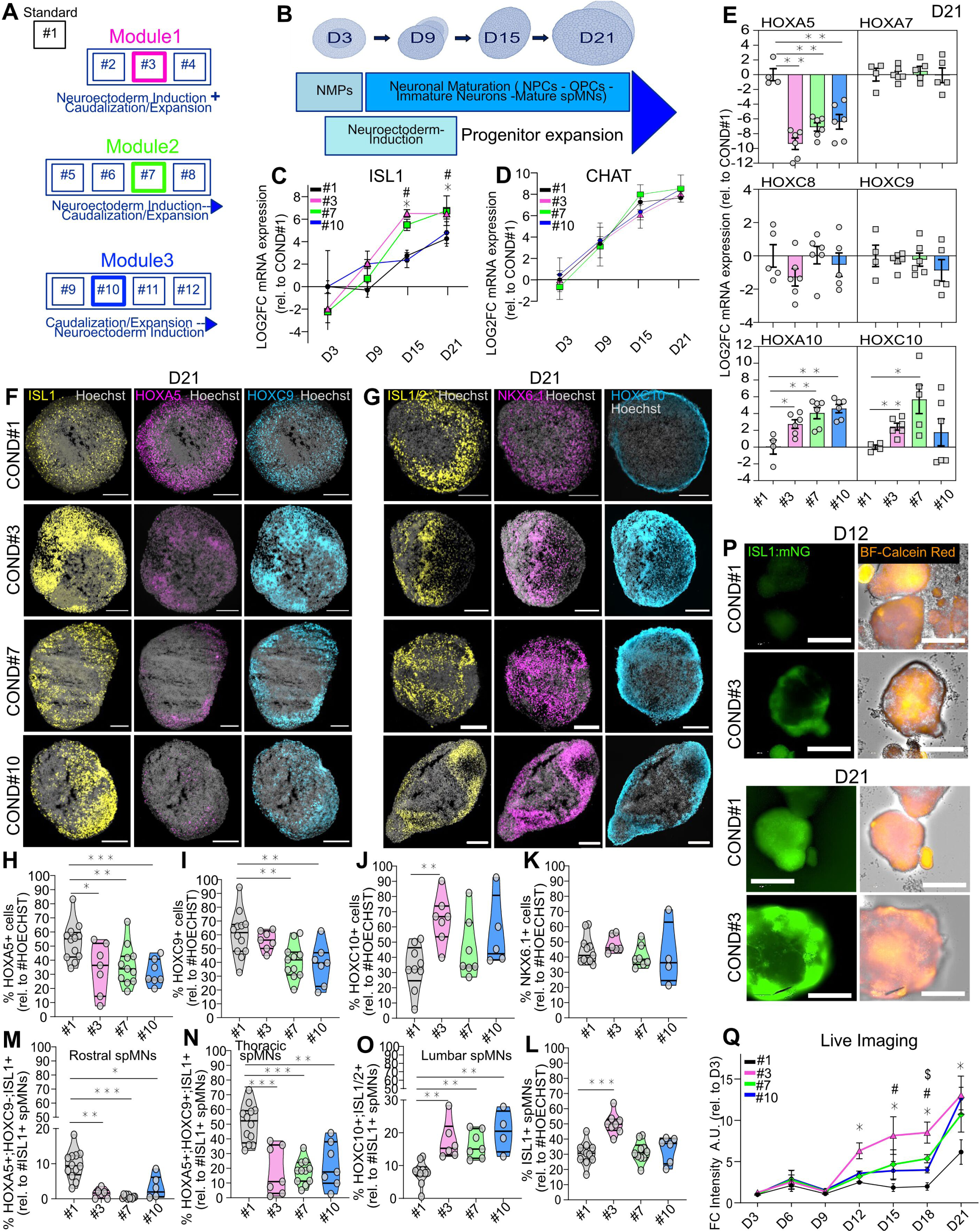
Neural induction during further caudalization of RA-primed NMPs generates lumbar spMNs. **A.** Patterning protocol screen design for guided differentiation of healthy hiPSC-derived and primed NMPs was conducted. The screen was divided into three modules, according to the sequence of neural induction and caudalization. **B.** SCOrgs derived from RA-primed NMPs were collected at several time points (day 3, day 9, day 15, day 21) to approximate the neurogenic timeline stage-dependently, and to phenotype neurogenesis rate and axial identity according to the protocol. **C.** qRT-PCR on several time points (see B), showing *ISL1* transcript levels. (One-way ANOVA with Fishers LSD or unparametric Mann-Whitney-U test. N=3 differentiations with n=3 technical replicates, each with 5-10 pooled spheres. Each symbol represents one sample). **D.** qRT-PCR on several time points (see B), plotting *CHAT* transcript levels. **E.** qRT-PCR on day 21 SCOs showing HOX expression. (Unparametric Mann-Whitney U test was used. N=2 differentiations with n=1-3 technical replicates, each with 5-10 pooled SCOs. Each dot represents a technical replicate). **F.** Representative images of cryosectioned day 21 SCOs immunostained against the spMN marker ISL1 with HOX genes for cervical (HOXA5, violet; thoracic (HOXC9, blue). Scale bar, 200µM. **G.** Representative images of cryosectioned day 21 SCOs immunostained against the spMN marker ISL1(2) with ventral domain marker NKX6.1(violet), and lumbar HOXC10 (blue). Scale bar, 200µM. **H.** Quantified immunostaining against cervical *HOXA5*. (Unparametric Kruskal-Wallis test with Uncorrected Dunn’s multiple comparison test. N=2-3 differentiations with n=1-4 SCOs per experiment. Each dot represents one organoid). **I.** Quantified immunostaining against thoracic HOXC9. **J.** Quantified immunostaining against lumbar HOXC10. **K.** Quantified immunostaining against ventral NKX6.1. **L.** Quantified immunostaining against spMN marker ISL1. **M.** Quantified co-immunostaining against HOXA5, HOXC9, ISL1 for cervical spMNs. **N.** Quantified co-immunostaining against HOXA5, HOXC9, ISL1 for thoracic spMNs. **O.** Quantified co-immunostaining against HOXC10, ISL1/2 for lumbar spMNs. **P.** Representative images from longitudinal high-content live imaging of SCOs derived from BJ- ISL1:P2A::mNG iPSC-Reporter Line at day 3 and day 21, shown for Condition#1 and #3. Cell culture was according to the novel differentiation protocols. Bright field was overlaid with Calcein AM (Orange) to allocate the Isl1-signal to individual SCOs. Scale bar, 2000µM. **Q.** ISL1-mNeonG fluorescence Intensity (A.U.) by area in the SCOs. (Two-way ANOVA with Fisher’s LSD Exact Test. N=2 experiments with n=4 wells per Corning P24 ULA plate. Each well contained between 10-50 SCOs. Each symbol represents the average of related mean fluorescence intensity per time point and protocol across all organoids). Graphics show mean+SEM, *p < 0.05, **p < 0.01, ***p < 0.001. *compares #1 with #3. $ compares #1 with #7. # compared #1 with #10.

Gene expression analysis from day 9 spheres showed that exposing the hiPSC-induced NMPs to CHIR+FGF2 for longer than the commonly applied 72h (Gouti et al. 2014, Lippmann et al. 2015, Faustino-Martins et al. 2020, Grass et al. 2024), resulted in higher levels of the caudal gene *FGF8* than when the NMPs were switched to neural induction signals earlier (protocols #2-12 vs protocol #1) (Fig. S1B). This increase in caudal specification did not compromise their neural differentiation potential, as *SOX2* and *NES* levels did not significantly change when compared to the standard protocol (#1), except for condition #5 (Fig. S1 C-D). The rostro-to-caudal shift observed in the new protocols was further validated in day 21 SCOs, where the expression of the rostral *HOX5* was markedly decreased compared to protocol #1, and the expression of more caudal *HOXs* (*HOXA10*, *HOXC10*) was significantly increased (Fig. S1E). In addition, SCOs generated through the new protocols showed an enhanced maturation profile, as 3 fold higher levels of *ISL1* (mature spMN marker) and up to 4 fold lower levels of *MNX1* (immature spMNs), 2-3 fold reduction of *OLIG2* expression (pMNs), *NKX6.1* (ventral spinal cord domain progenitors) or *DCX* (newly differentiated neurons) indicated (Fig. S1 F). Immunostaining analysis validated the increased presence of caudal HOX^+^ cells in detriment of rostral ones (Fig. S1 G-I) in the new protocols compared to the standard #1. No clear difference in the number of OLIG2^+^ progenitors or NeuN^+^ neurons was detected across the protocols (Fig. S1 J-L), and interestingly, protocol #3 showed a 50% increase in the number of ISL1^+^ spMN (Fig. S1 M).

While all the new patterning protocols showed a higher caudalization potential than the standard one, no other major differences among them were detected. Thus, we selected one paradigm from each of the modules (#3, #7, and #10) to perform a longitudinal transcriptomic analysis along the SCO development in comparison to the standard protocol (#1) and test whether the new methods accurately recapitulated the temporal sequence of in vivo developmental processes (Sagner et al 2021). In all conditions, transcript levels of early neural development (*SOX2*, *PAX6*) appropriately decreased as development progressed (Fig. S2 A-B), while markers of more advanced stages during spinal cord development (*NKX6.1* and *SYN1*) showed an expected simultaneous increase (Fig. S2 C-D). Interestingly, SCOs generated from the novel protocols showed a more rapid decline in *OLIG2* levels from day 9 to day 21, suggesting a faster progression away from progenitor states and towards more differentiated neural fates (Fig. S2E). Furthermore, while analogous levels of the mature spMN marker *CHAT* were detected throughout SCO development across the three selected protocols, and similar to the standard protocol #1, spMN *ISL1* transcript levels at day 21 were significantly elevated in protocols #3 and #7 (Fig. 3C-D). This increase in *ISL1* expression points again to enhanced spMN formation under these conditions, in alignment with our qPCR and immunostaining results from the 12 protocol screen study (Fig. S2F-G). qPCR-analysis of RC *HOX* expression showed a marked decrease of rostral *HOXA5* transcript for the three new protocols compared to standard #1, no significant changes for the thoracic *HOXA7*, *HOXC8* or *HOXC9* and a marked increase in lumbar *HOXA10* and *HOXC10* transcripts, especially for protocols #3 and 7 (Fig. 3E). To validate our gene expression data, we next carried out extensive immunostaining analysis of day 21 SCOs generated through the three selected protocols. As expected with the addition of extended NMP maintenance state, the novel protocol paradigms induced an axial identity shift in the SCOs. Compared to the SCOs generated following protocol #1, the new SCOs showed 10-15% less HOXA5^+^ cervical cells (Fig. 3F, H) and a 20% less thoracic HOXC9^+^ cells (Fig. 3F, G). Conversely, ∼30% increase of lumbar cells with nuclear HOXC10^+^ was found for the selected new protocols, and significantly for protocol #3 (Fig. 3G, J). No differences in the number of cells positive for the ventral domain marker NKX6.1 were detected in immunostaining (Fig. 3G, K), albeit we noticed significant differences in *NKX6.1* gene expression (Fig. S2C). In addition, protocol #3 again showed a significant increase in the percentage of ISL1^+^ spMNs compared to other protocols (Fig. 3F, G, L, P, Q). To deepen our analysis of spMN axial identity in the SCOs generated by the selected new protocols, we quantified the percentage of ISL1^+^ spMNs with rostral, thoracic or lumbar HOX identity. The new protocols resulted in a drastic decrease in ISL1^+^;HOXA5^+^;HOXC9^-^ cervical spMNs (Fig. 3M), and a mild decrease in ISL1^+^;HOXA5^+^;HOXC9^+^ thoracic spMNs (Fig. 3N), accompanied by a moderate increase in ISL1^+^;HOXC10^+^ lumbar spMNs (Fig. 3O).

Lastly, to further validate the efficiency of our new protocols at producing spMNs, we generated a reporter hiPSC line where the green fluorescent protein mNeonGreen (mNG) was knocked-in downstream of the endogenous *ISL1* locus (Fig. S2H-I). Using this reporter line, we live-tracked spMN differentiation during the maturation of SCOs generated according to the 12 protocols. Using this approach, the enhanced generation of spMNs by the new protocols compared to the standard protocol #1 was particularly evident for protocol #3 (Fig. 3P-Q and S2J). In summary, SCOs generated using our new experimental differentiation modules exhibited greater spMN RC diversity without reducing overall spMN specification, compared to the commonly used NSC-based differentiation condition #1 that primarily yields spMNs with a rostral identity. Overall, protocol #3 consistently produced the highest quantity of ISL1^+^ spMNs, as confirmed by qRT-PCR, immunostaining and live imaging. This protocol also showed a shift from cervical to lumbar *HOX* gene expression compared to the standard protocol. Together, we conclude that RA-primed NMPs, when subjected to differentiation protocol #3, efficiently generate spMNs with a broader RC molecular identity than the standard method. However, this protocol generated only ∼2.5% of rostral spMNs (HOXA5+:HOXC9-:ISL1+). This highlights the need to achieve a balanced axial heterogeneity of spMNs within the same organoid by means other than modifying the patterning protocol.

### Pre-mixed NSCs/RA-primed NMPs generate SCOs comprised of progenitors with rostral and caudal neural identities

While the commonly used NSC-based SCO protocols primarily produce hindbrain and cervical spMNs (>80%), our selected protocol paradigm #3, starting from RA-primed NMPs yielded an overall increased spMN population with predominantly thoracic and lumbar identities. Although this new protocol does not completely halt the specification of rostral spMNs, our aim was to develop an improved approach able to generate a balanced RC spMN diversity for a more physiologically relevant in vitro model to study spinal cord development and disease conditions. To achieve a greater axial heterogeneity within a single organoid, we therefore explored whether using a premix pool of RA-primed NMPs and NSCs as a starting stem cell population, rather than only RA-primed NMPs, would achieve that goal (Fig. 4A). As different predisposition to differentiate into rostral or caudal fates between different hiPSC lines has been reported (Kim et al. 2024), we generated NSCs and RA-primed NMPs under adherent conditions from multiple healthy lines (BJ-siPS, 1016A, KOLFC1, 18A, CRTD1) and then used them in an even mixture to generate stem cell aggregates. Our adherently generated NSCs were clearly immunoreactive for panneuronal SOX1 and the stem cell marker REST (Kojima et al. 2001, Hwang et Zukin 2018), whereas NMPs were positive for caudal CDX2 and REST (Fig. S3A). As control conditions for our SCO patterning, either only RA-primed NMPs or only NSCs were used as starting stem cell populations. We characterized the resulting SCOs by longitudinally evaluating the expression of key genes governing spinal cord development and spMN specification during the SCO development by immunostaining and qRT-PCR. Notably, "NSC-NMP Mix" aggregates assembled robustly and uniformly (Fig. 4B) and, at day 0, showed immunoreactivity for approximately equal levels of the mesodermalal lineage marker TBXT and the neural lineage marker PAX6, with a slightly lower amount of SOX2^+^ NSCs (Fig. S3B-C). In contrast, NMP spheres showed TBXT, SOX2 and CDX2 expression without PAX6, while NSC spheres showed SOX2, PAX6 and neural NEUROD1 expression without TBXT or CDX2 (Fig. S3B-C). The NMP spheres also showed highest expression of other caudal progenitor and NMP marker genes, such as *NT5E* and *HOXC9* (Fig. S3D). We observed that in the mixed SCOs, both progenitor cell populations segregated into SOX2^+^:CDX2^-^ and SOX2^-^:CDX2^+^ areas (Fig. 4C).

**Figure 4.**
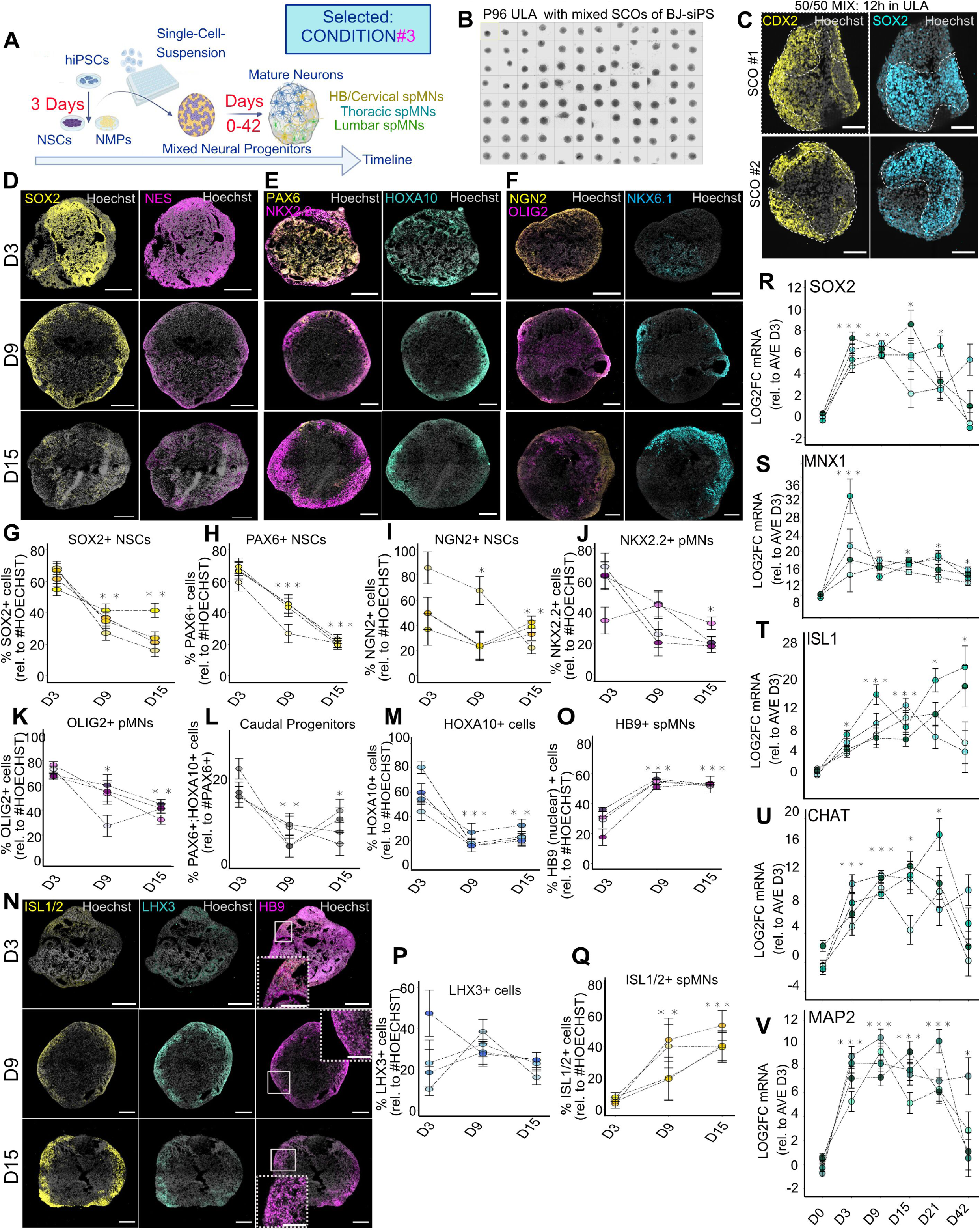
An even NMP/NSC mix as a starting stem cell population to generate SCOs results in a balanced specification of rostral and caudal NPCs. **A.** Schematic representation of the experimental design followed to generate SCOs. **B.** Bright field image of one representative 96-well ULA plate showing spinal spheres 12h after stem cell aggregate formation. **C.** Immunostaining against caudal CDX2 (yellow) and neural SOX2 (blue) on 50% NSC:50% NMP aggregates at day 0, generated from the BJ-siPS hiPSC line, segregation is shown by the dotted white line. Scale bar, 200µM. **D.** Immunostaining against the NSC markers SOX2 (yellow) and NESTIN (violet) on day 3, 9 and 15 SCOs made from a BJ-siPS 50% NSC:50% NMP mix subjected to Protocol #3. Scale bar, 200µM. **E.** Immunostaining against NSC PAX6 (yellow), caudal progenitor NKX2.2 (violet) and lumbar HOXA10 (green) markers on day 3, 9 and 15 SCOs made from a BJ-siPS 50% NSC:50% NMP mix subjected to Protocol #3. Scale bar, 200µM. **F.** Immunostaining against neural progenitor NGN2 (yellow), pMN OLIG2 (violet) and ventral spinal cord domain NKX6.1 (blue) markers on day 3, 9 and 15 SCOs made from a BJ-siPS 50% NSC:50% NMP mix subjected to Protocol #3. Scale bar, 200µM. **G.** Quantification of the percentage of SOX2+ NSCs on BJ-siPS 50:50 SCOs subjected to Protocol #3 over time. (Friedman Test with Dunn’s posthoc test. Each dot represents the mean from all organoids of one differentiation batch. Time-points from a batch are connected by a line and have the same colour. N=4 differentiations with n=3-8 SCOs per experiment). **H.** Quantification of the percentage of PAX6+ NSCs on SCOs generated as in (G). **I.** Quantification of percentage of counted NGN2+ NSCs on SCOs generated as in (G). **J.** Quantification of percentage of NX2.2+ pMNs vental-caudal progenitors on SCOs generated as in (G). **K.** Quantification of percentage of counted OLIG2+ pMNs on SCOs generated as in (G). **L.** Quantification of percentage of PAX6+:HOXA10+ caudal progenitors on SCOs generated as in (G). **M.** Quantification of percentage of HOXA10+ cells on SCOs generated as in (G). **N.** Immunostaining against mature spMN marker ISL1/2 (yellow), immature spMN marker HB9 (violet) and LIM domain factor LHX3 (green) on day 3, 9 and 15 SCOs made from a BJ-siPS 50% NSC:50% NMP mix. Scale bar, 200µM. **O.** Quantification of percentage of HB9+ spMNs on SCOs generated as in as in (N). **P.** Quantification of percentage of LHX3+ cells on SCOs generated as in as in (N). **Q.** Quantification of percentage of ISL1/2+ spMNs on SCOs generated as in as in (N). **R.** Longitudinal qRT-PCR analysis showing mean LOG2FC expression of *SOX2* gene from day 0 to day 42 in BJ-siPS 50% NSC:50% NMP mix SCOs subjected to Protocol #3. (Friedman Test with Eisinga’s correction. Each dot represents one sample from 3-8 pooled SCOs, N=4 experiments. Time-points from a batch are connected by a line and have the same colour). **S.** Longitudinal qRT-PCR showing mean LOG2FC of *HB9* transcript on SCOs generated as in (S). **T.** Longitudinal qRT-PCR showing mean LOG2FC of *ISL1* transcript on SCOs generated as in (S). **U.** Longitudinal qRT-PCR showing mean LOG2FC of *ChAT* transcript on SCOs generated as in (S). **V.** Longitudinal qRT-PCR showing mean LOG2FC of *MAP2* transcript. For all graphics, mean± SEM or LOG2FC ± SEM is shown, *p < 0.05, **p < 0.01, ***p < 0.001.

Next, we investigated whether the SCOs generated from a NSC:RA-primed NMP mixture and subjected to our new differentiation protocol #3 matured normally according to the embryonic development of the spinal cord (Sagner et al 2019). As expected, the number of NSCs (SOX2^+^, PAX6^+^) and NPCs (NGN2^+^, NKX2.2^+^) decreased from 70% to 30% from day 3 to day 15 (Fig. 4D-J), similarly to the number of spMN progenitors (OLIG2^+^) (Fig. 4F, K). We also observed that the number of PAX6^+^;HOXA10^+^ caudal progenitors and the total number of HOXA10+ cells decreased as the SCOs developed (Fig. 4E, L and 4E, M, respectively).

We next examined the emergence of spMN specific markers. Consistent with the literature, as the ventral progenitors in the SCOs acquired a spMN identity, the number of cells showing a nuclear HB9 staining, indicative of early-born spMNs (Letchuman et al. 2022), increased from day 3 to day 9-15 (Fig. 4N-O). The quantification of the number of LHX3^+^ cells, a transcription factor that together with ISL1 mediates the expression of genes critical for medial motor column spMN specification and maturation (Erb et al. 2017; Smith et al. 2020) increased from day 3 to day 9 (Fig. 4N, P), as well as the number of ISL1/2^+^ spMNs (Fig. 4N, Q), indicating the differentiation of ventral progenitors into specific spMN subtypes. We validated our immunostaining-based results via longitudinal expression analysis. As expected, the developmental gene expression patterns from day 0 to day 42 SCOs generally matched our cell counts. *SOX2* peaked at day 3 and sharply declined thereafter (Fig. 4R) and the early spMN marker HB9 was highest at day 3 (Fig. 4S). It is important to note that HB9 is expressed in late, differentiating NMPs, as well as in immature spMNs, which could explain the 30 fold increase in expression from day 0 to day 3 (Verrier et al. 2018). Moreover, the expression of more mature neuronal markers *MAP2*, *ISL1* and *CHAT* gradually increased, reaching a plateau past day 21 (Fig. 4T-V). All these results were recapitulated using two additional healthy hiPSC lines (Fig. S4). Overall, these results demonstrate that SCOs generated using an even mix of NSCs/RA-primed NMPs and subjected to the protocol #3 fairly recapitulate in vivo spinal cord developmental differentiation processes (Sagner et al. 2018, Sagner et al. 2019).

### Mature SCOs derived from the mixed stem cell pool are composed by a balanced axial spMNs diversity

Once we validated that the SCOs produced from the pre-mixed pool of adherently-generated NSCs and RA-primed NMPs and patterned according to protocol #3 contained neural progenitors with rostral and caudal differentiation potentials, we next decided to systematically quantify the number of hindbrain, cervical, thoracic and lumbar cells and spMNs derived from those progenitors in day 21 SCOs. We analysed SCOs generated from several healthy lines using the three different stem cell aggregate compositions (100% NSCs, 100% RA-primed NMPs and 50%:50%) to determine whether the even mixture of two different stem cell populations resulted in a larger axial heterogeneity and higher abundance of spMN compared to the other two approaches. First, we observed that the relative proportion of ISL1^+^ (or ISL1^+^/2^+^) spMNs remained consistent (∼40%) across conditions and lines (Fig. 5A-C and S5A-K). Next, SCOs derived from the "50:50 mix" condition yielded 24% cervical-brachial (HOXB4^+^;HOXC10^-^, HOXA5^+^;HOXC9^-^ or HOXC6^+^;HOXA10^-^) (Fig. 5A-F, K and S3E-J), 48% thoracic (HOXA5^-^;HOXC9^+^) (Fig. 5B, G, K and S3K-L), and 28% lumbar (HOXC6^+^;HOXA10^+^ or HOXB4^+^;HOXC10^+^) (Fig. 5A, C, H-I, K and S3M-P) spMNs.

**Figure 5.**
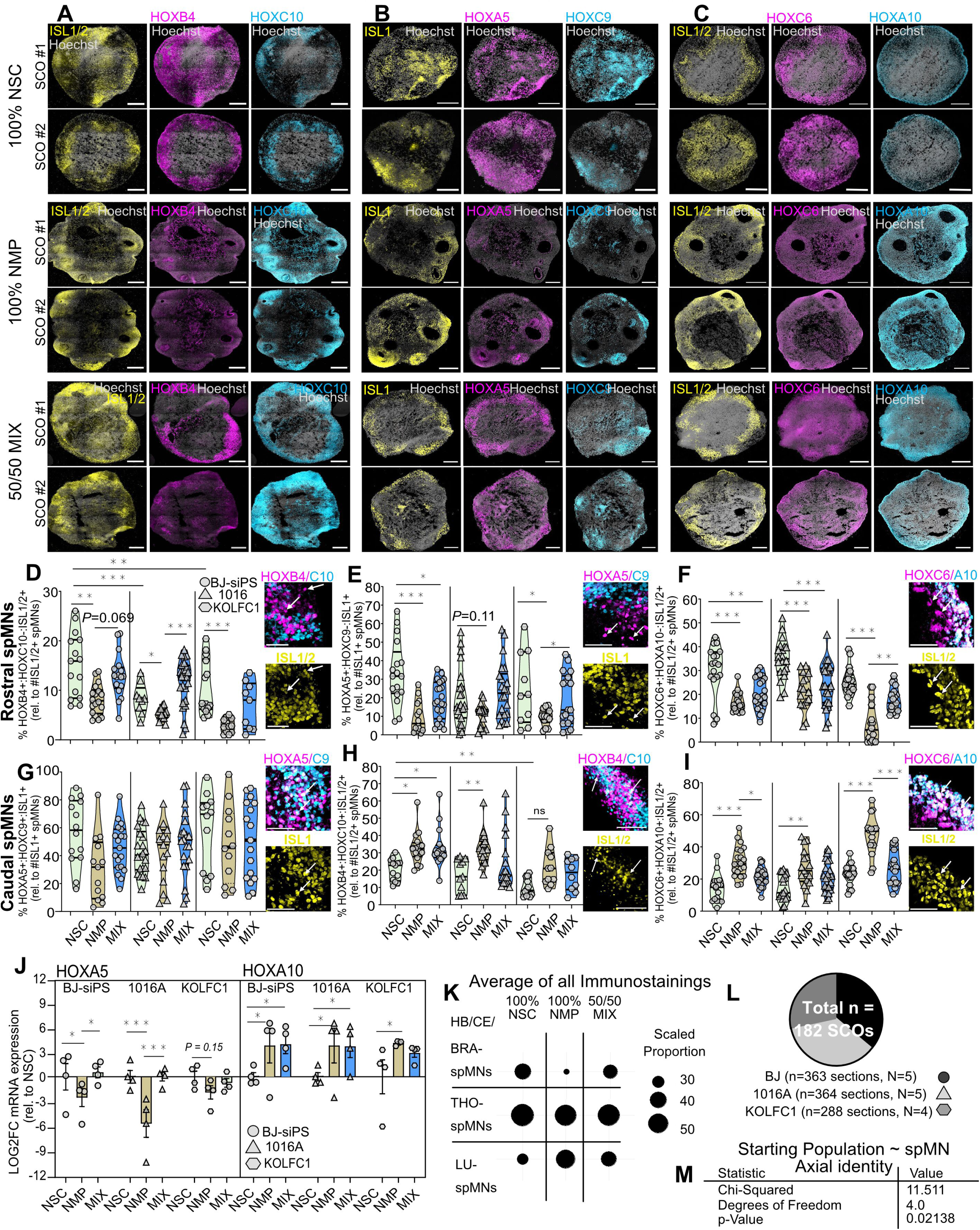
SCOs derived from the mixed stem cell populations are composed by spMNs with high axial heterogeneity (CASCOs) **A.** Immunostaining against ISL1/2 (yellow), HOXB4 (violet) and HOXC10 (blue) on day 21 BJ-sPS SCOs generated according to three differentiation conditions: 100% NSCs, 100% NMPs, and 50% NSCs:50% NMPs. Scale bar, 200µM. **B.** Immunostaining against ISL1 (yellow), HOXA5 (violet) and HOXC9 (blue) on day 21 SCOs as in (A). **C.** Immunostaining against ISL1/2 (yellow), HOXC6 (violet) and HOXA10 (blue) on day 21 SCOs as in (A). **D.** Quantified co-immunostainings against ISL1/2+; HOXB4+; HOXC10- on day 21 SCOs generated according to three differentiation conditions from three healthy cell lines (BJ-siPS, 1016A, KOLFC1). Hereby, we show hindbrain spMNs. (Violin plots show median and interquartile range. N=4-5 differentiations, n=3-6 SCOs per experiment. Each dot represents one organoid, and is the average cell ratio from n=1-3 imaged cryosections per organoid. Two- way ANOVA with Tukey-Kramer post hoc test and heteroscedastic adjustment). Showing Zooms of CASOs with white arrows indicating two examples for the axial spMN population that was identified. Scale bar in Zooms, 50µM. **E.** Quantified co-immunostainings showing the percentage of cervical spMNs (ISL1+; HOXA5+; HOXC9-) as in (D). **F.** Quantified co-immunostainings showing the percentage of brachial spMNs (ISL1/2+ HOXC6+, HOXA10-) as in (D) **G.** Quantified co-immunostainings showing the percentage of thoracic spMNs (ISL1+; HOXA5+; HOXC9+) as in (D). **H.** Quantified co-immunostainings showing the percentage of lumbar spMNs (ISL1/2+; HOXB4+; HOXC10+) as in (D). **I.** Quantified co-immunostainings showing the percentage of lumbar spMNs (ISL1/2+ HOXC6+, HOXA10+) as in (D). **J.** qRT-PCR analysis on day 21 SCOs generated as in (A) showing *HOXA5* and *HOXA10* transcript levels. (Unparametric Mann-Whitney-U test. Each dot corresponds to one sample obtained from an independent differentiation batch (N=4, n=1) and was generated by pooling 3 SCOs). **K.** Average scaled percentage of spMNs with specific RC anatomical HOX identity assigned through co-immunostaining of ISL1 and HOX genes. Data from all SCOs analysed from all experiments performed as part of this figure (see L) and cell lines is compiled. **L.** Pie-Chart with overview of all analysed cryosections from all SCOs as part of this figure (biological replicates). **M.** Table showing dependence and statistical testing of the factors “spMN patterning” against “starting population of stem cells”. For all graphics, LOG2FC, mean±SEM are shown, *p < 0.05, **p < 0.01, ***p < 0.001.

These data fit with qRT-PCR analysis to assess the relative abundance of RC *HOX* mRNA transcripts across the three organoid conditions. NMP-derived SCOs showed a marked decrease in rostral *HOX* gene expression relative to the NSC and the mix SCOs. Conversely, NSC-derived SCOs displayed significantly lower caudal *HOX* gene expression levels than the NMP and the mix SCOs (Fig. 5J and S3Q). Additionally, we observed that spMN subpopulations with different axial identities self-arranged in a regionalized manner within the SCOs segregating from each other (Fig. 5 A-C and S5L-N). Overall, we observed a more balanced proportion of RC spMN identities in NSC:NMP SCOs (25:47:27) than in the ones generated solely from NSCs or NMPs (30:51:19 and 15:47:38, respectively (Fig. 5K). This suggests that the relative proportion of RC spMN subtypes generated seemed to depend on the starting population of stem cells used for SCO formation. Importantly, statistical analysis based on 1015 imaged SCO cryosections from a total of 182 processed SCOs for ISL1/2 and HOX immunostaining (Fig. 5L) showed statistical dependence of the “spMN Axial Subtype” factor on the “Starting Cell Population (NSC, NMP, MIX)” factor (Fig. 5M).

In summary, while the specification of ISL1/2^+^ spMNs did not differ across the three conditions and is therefore not dependent on the starting stem cell population, at least when subjected to our protocol condition #3, using an even mix of NSCs and RA-primed NMPs as the starting stem cell pool consistently enhanced the axial heterogeneity of the resulting spMNs. Together, this new organoid patterning protocol, that we name comprehensive axial spinal cord organoid (CASCO), constitutes a highly robust and efficient method for directional differentiation of neural progenitor cells into a wide variety of axial spMN identities.

### Inducible vasculature and skeletal muscle are beneficial for neural maturation in CASCOs

Most organoid models, particularly SCOs, suffer from the absence of vascular networks, which restricts oxygen and nutrient delivery to the innermost regions of the organoid. In addition, a crosstalk between spMNs and endothelial cells has been reported necessary for a correct spMN development (Vieira et al. 2022). Similarly, when modelling neuromuscular development and disease, integrating the spMN cell targets, muscle cells, into the organoids is critical to recapitulate the motor unit. As for endothelial cells, muscle fibers release growth factors that contribute to spMN development and maturation (Saini et al. 2021). To increase the resemblance of our new CASCO protocol to the in vivo environment, we generated CASCOs comprising a functional endothelial cell network and skeletal muscle cells intertwined with the neural tissue (Fig. 6A). For this, we used a transposase-based approach to engineer hiPSCs carrying an inducible *ETV2* overexpressing construct that, upon doxycycline treatment, induces hiPSCs differentiation to endothelial cell lineage (Cakir et al. 2019, Ng et al. 2021). SCOs generated from this edited line, after being exposed to dox during the entire duration of the differentiation protocol and immunostained against the mature vascular marker CD31, showed tubular-like structures resembling endothelial micro-vessels that were integrated within the organoid tissue (Fig. 6B). Once we validated the efficiency of this line to differentiate into endothelial CD31^+^ cells and their self-organization in the organoids, we generated CASCOs from unedited and iETV2 hiPSC lines in an 80:20 cell ratio and studied several indicators of neural maturation commonly used in the organoid field (Hergenreder et al. 2024). High-resolution imaging of lineage-mixed CASCO cryosections on day 42 showed a significant increase in the number of Synaptophysin^+^ (presynaptic marker) and PSD95^+^ (post-synaptic) puncta compared to CASCOs only comprised of neural tissue (Fig. 6C-F, G-H).The levels of the trophic factor BDNF, associated with synaptic plasticity and activity-dependent neuronal differentiation and maturation, were also significantly increased in the CASCOs+iETV2 condition (Fig. 6E, I). Importantly, the number of mature spMNs stained with CHAT antibody was not reduced in the CASCOs+iETV2 organoids, despite these contained 20% fewer cells permissive to become spMNs due to the presence of dox-inducible endothelial cells (Fig. 6E-F, J). These results indicate that introducing endothelial cells in our CASCOs, capable of self-assembling into vasculature-like structures, enhances the maturation of the neurons present in the mixed-lineage SCO.

**Figure 6.**
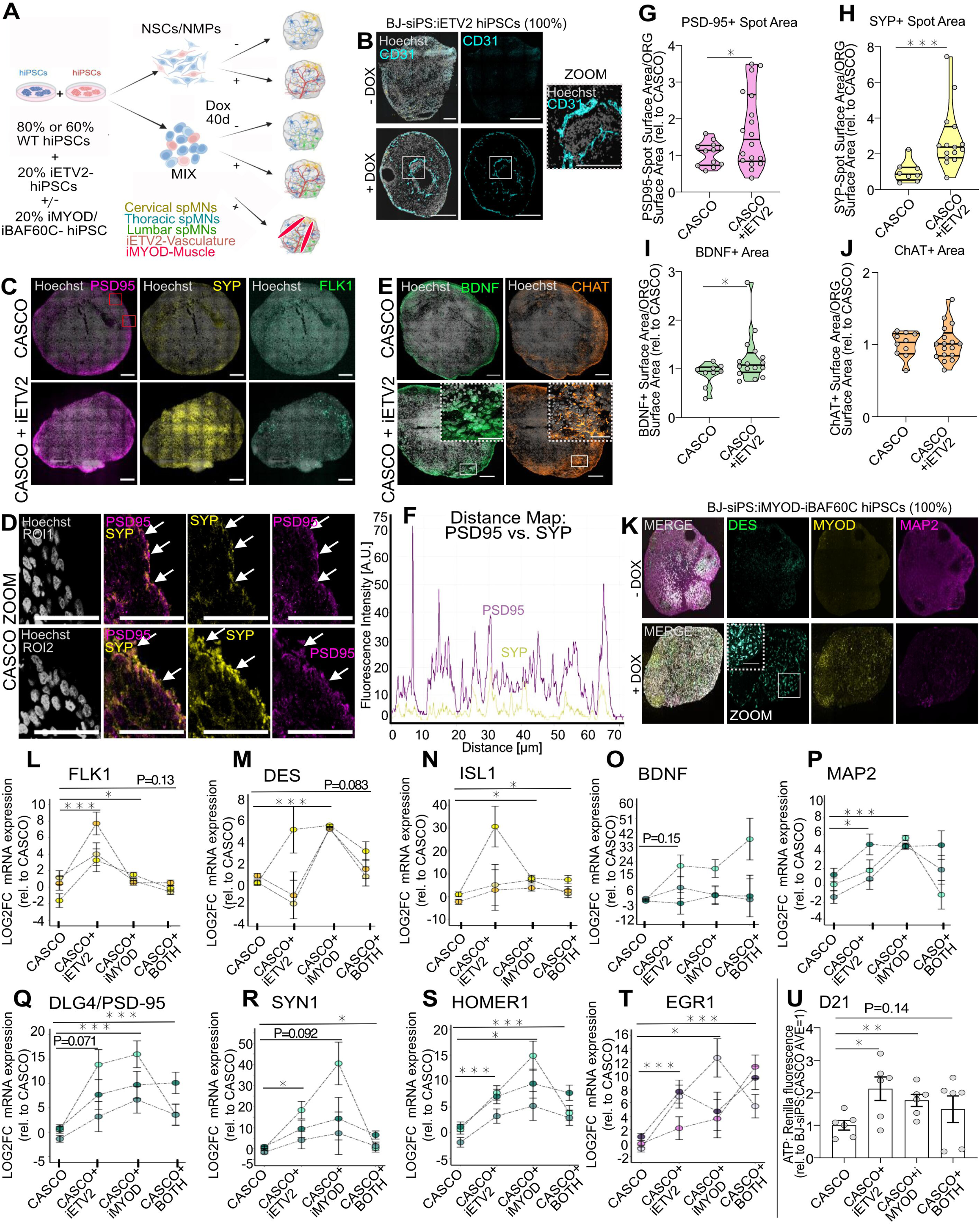
Inducible vasculature and skeletal muscle are beneficial for neural maturation in CASCOs. **A.** Scheme illustrating the approach followed to generate complex CASCOs from either only unedited hiPSCs (BJ-siPS line) or a mixture of unedited hiPSCs and hiPSCs expressing doxycycline-inducible ETV2 (transcription factor governing vascular endothelial cell specification) and/or hiPSCs expressing doxycycline-inducible MYOD/BAF60C (transcription factors promoting myoblast specification) (inducible tri-lineage organoids). **B.** Representative immunostaining and magnification (right) against the mature vascular surface marker CD31 (blue) on day 28 complex CASCOs generated from a 100%:iETV2 hiPSCs without and with doxycycline treatment. Note that the CD31+ cells are self-organized in tube–like structures. Scale bar, 200µM. Scale bar of Zoom, 50µM. **C.** Representative immunostaining against the synaptic markers PSD-95 (violet) and Synaptophysin (yellow) and the endothelial marker FLK1 (green) on day 42 CASCOs and CASCO+iETV2 showing synaptic punctae and endothelial cells. Scale bar, 200µM. **D.** Magnification of immunostained organoid cryosections using confocal AiryScan showing synaptic staining as in (D). Scale bar, 100µM **E.** Representative immunofluorescence staining against the neurotrophin BDNF (green) and the mature spMN marker CHAT (orange) on day 42 CASCOs and CASCO+iETV2. Scale bar, 200µM. Scale bar of Zoom, 50µM. **F.** Distance Map showing the physical separation of PSD-95 (violet) and Synaptophysin (yellow) staining signal, with synaptic punctae following a similar distance-peak pattern. **G.** Quantification of organoid cryosection area occupied by PSD95 puncta on day 42 BJ-siPS CASCOs and CASCO+iETV2. (One-sided, unpaired t-test with Welch’s correction. N=4 independent CASCO-differentiation batches, n=1-3 organoids/experiment. Each dot corresponds to one organoid). See D. **H.** Quantification of organoid cryosection area occupied by SYP puncta as in (D). **I.** Quantification of organoid cryosection area occupied by BDNF puncta as in (E). **J.** Quantification of organoid cryosection area occupied CHAT-staining as in (E). **K.** Representative immunostaining and magnification (right) against the muscle markers MYOD (yellow) and DESMIN (green) and the pan-neural MAP2 (violet) on day 21 CASCOs generated from a 100% iMYOD/BAF60C hiPSCs in the absence or presence of doxycycline. Scale bar, 200µM. **L.** qRT-PCR analysis showing endothelial *FLK1* mRNA expression as in (L). from day 42 CASCOs or complex CASCOs+iETV2, CASCOs+iMYOD/BAF60C or CASCOs+iETV2+ iMYOD/BAF60C organoids (Friedman Test with Eisinga’s correction. Each dot corresponds to one sample obtained from an independent differentiation batch (N=3, n=1) and was generated by pooling 3 day 42 SCOs). **M.** qRT-PCR analysis showing muscle marker *DES* mRNA as in (L). **N.** qRT-PCR analysis showing spMN marker *ISL1* mRNA expression as in (L). **O.** qRT-PCR analysis showing neurotrophin *BDNF* mRNA expression as in (L). **P.** qRT-PCR analysis showing pan-neural marker *MAP2* mRNA expression as in (L). **Q.** qRT-PCR analysis showing *DlG4* mRNA expression (coding for PSD95) as in (L). **R.** qRT-PCR analysis showing synaptic marker *SYN1* mRNA expression as in (L). **S.** qRT-PCR analysis showing synaptic marker *HOMER1* mRNA expression as in (L). **T.** qRT-PCR analysis showing immediate early gene *EGR1* mRNA expression as in (L). **U.** Renilla luminescence quantification (CellTiter-Glo Luminescent assay) on day 21 CASCOs or complex CASCOs+iETV2, CASCOs+iMYOD/BAF60C or CASCOs+iETV2+ iMYOD/BAF60C organoids (Non-parametric Kruskal-Wallis and uncorrected Dunn’s test. N=4 independent differentiations, n=1-2 SCOs per experiment. Each symbol represents one technical. For all graphics, mean±SEM is shown, *p < 0.05, **p < 0.01, ***p < 0.001.

Several studies have developed 2D co-cultures of hiPSC-derived spMNs and skeletal muscle cells or 3D neuromuscular organoids using various techniques to model and study neuromuscular junction formation in healthy and disease conditions (Demestre et al. 2015, Lenzi et al. 2016, Tiago et al. 2021, Urzi et al. 2024). Often these approaches include the addition of lineage-dependent cues and/or the supplementation of small molecule cocktails to induce the specification of the desired cell type and enhance organoid maturation (Faustino-Martins et al. 2020, Hergenreder et al. 2024). Using an analogous approach to the one undertaken for our CASCO+iETV2 system, we edited the same hiPSC line to introduce dox-inducible MYOD1 and BAF60C2 transcription factors, known to regulate muscle differentiation (Weintraub et al. 1991, Meng et al. 2018, Tiago et al. 2021). Myospheres generated using this edited hiPSC line and cultured under dox treatment broadly contained DESMIN^+^ and MyoD^+^ skeletal muscle cells (Fig. 6K). Next, we generated CASCOs containing both inducible lineage-specific hiPSC lines in a 60:20:20 ratio (WT:iETV2:iMYOD+BAF60C) and compared the expression levels of endothelial, skeletal muscle, spMN/neuron and synaptic/activity markers across the three types of complex CASCOs (CASCO+iETV2, CASCO+iMYOD+BAF60C and CASCO+iETV2:iMYOD+BAF60C) relative to the control CASCOs only comprised of neural tissue at a mature developmental stage (day 42). We first quantified the mRNA levels of lineage-confirming genes in the organoids. As expected, the expression levels of the endothelial marker *FLK1* were highest in the CASCO+iETV2 condition (Fig. 6L) and the skeletal muscle marker *DESMIN* expression was highest in the CASCO+iMYOD+BAF60C organoids (Fig. 6M). Interestingly, despite being comprised by a lower percentage of cells committing to neural lineage (20% and 40% less in the bi- and tri-lineage organoids, respectively), the complex CASCOs containing either endothelial, muscle cells or both, showed increased *ISL1* and *MAP2* levels compared to the control CASCOs (Fig. 6N, P). Moreover, we observed an increase in the expression of *BDNF* (Fig. 6O) and the synaptic *DLG4* (*PSD95*), *SYN1* and *HOMER1* in the complex organoids (Fig. 6 Q-S), and enhanced expression of the immediate early gene *EGR1* (Duclot et Kabbaj 2017) compared to control CASCOs (Fig. 6T). These results suggest that the presence of endothelial and skeletal muscle cells in the organoids boosts the maturation of the existing neurons. Lastly, we detected that the presence of these non-neural cell lineages was beneficial for the overall organoid health, as measurement of ATP levels, a proxy for cell viability and metabolic activity, showed higher values in the multi-lineage CASCOs than in the control ones (Fig. 6U). In conclusion, adding both a vasculature-like network and skeletal muscle cells developmentally intertwined with the neural tissue considerably improved the maturation and overall health state of the CASCOs.

## Discussion

In recent years, several studies using NGS approaches have described the cellular composition of the developing and adult human spinal cord, providing an invaluable tool to identify specific spMN subtypes based on precise transcriptomic signatures (Rayon et al. 2021; Yadav et al. 2023). Numerous protocols for generating SCOs from hiPSCs are also now available (Xu et al. 2023; Whye et al. 2023; Gribaudo et al. 2023; Mouilleau et al. 2021; Ogura et al. 2018; Faustino-Martins et al. 2020; Grass et al. 2024). However, currently, no methodology enables the production of a high amount and diversity of rostro-caudal spMNs within the same organoid. Such an approach would be instrumental for exploring the developmental cues governing human axial spMN specification, like gene regulatory networks (Dasen et al. 2005, Lacombe et al. 2013), the sequence of key transcription factor activation during spMN development (Novitch et al. 2001, Sagner et al. 2019) or the acquisition of columnar identiy (Sockanathan et Jessell 1998, Baek et al. 2017). Furthermore, in motor neuron diseases like ALS or SMA, spMNs corresponding to specific spinal segments show distintct vulnerabilities to the disease (Simon et al. 2017; Nizzardo et al. 2020), however the underlying mechanisms remain obscure (Kampmann 2024). An organoid model containing both resistant and vulnerable axial spMN populations would facilitate examining how specific stressors impact both subtypes, whether they cope differently with an external insult or when they acquire their divergent vulnerability nature. While recent studies using direct programming of PSCs into different MN types have started to elucidate distinct neuronal sensitivities to disease (An et al. 2019), this approach drives differentiation in an extremely fast manner that skips neurodevelopmental intermediates, which could hamper the discovery of mechanisms triggering the pathology. Additionally, this approach results in homogenous populations of cells, which lacks the rich heterogeneity of the nervous system. Historically, the protocols used to produce spMNs from PSCs gave rise primarily to neurons with rostral identity. While in recent years other protocols for patterning SCOs toward more lumbar fates have been developed, these approaches face limitations, including a reduced number of mature spMNs, limited spMN heterogeneity or a complete loss of spMN subpopulations (Andersen et al. 2020; Ogura et al. 2018; Hor et al. 2018; Gribaudo et al. 2023). In this study, we aimed to generate a human SCO model comprising spMNs with molecular diversity across the entire anatomical axis of the spinal cord. By refining the induction of more caudal fates from RA- primed NMPs simultaneously patterned with rostral NSCs, we achieved a near-physiological anatomical specification of spMNs in our CASCO model (Fig. 5K), which are equivalent to the spMN profiles measured via scRNAseq along the spinal segments in the developing human spinal cord (Zhang et al. 2021). Furthermore, the inclusion of endothelial-like cells and skeletal muscle enhanced the physiological relevance and neural maturation of the model.

As the embryo undergoes axial extension, neural tube development occurs in two phases of neurulation that rely on two distinct progenitor populations, NSCs and NMPs, which contribute to the development of the CNS (Schoenwolf et Smith 1990, Shaker et al. 2021). Historically, most SCO protocols aimed at mimicking the formation of the rostral neuraxis. These protocols rely on the immediate induction of PSCs into NSCs, the first stem cell population contributing to the formation of the mammalian spinal cord. This induction is typically achieved through BMP pathway inhibition (Chambers et al. 2009). NSCs patterned in this way are fixed in their axial identity and therefore unable to generate more caudal neural structures (Lippmann et al. 2015). While these protocols generate a high percentage of spMNs, these have mainly a rostral identity (Hor et al. 2018), and thus these organoids have severely limited axial heterogeneity, which we corroborated via scRNAseq analysis. The second stem cell source and the one most recently used to generate SCOs, NMPs, is a bipotent cell population responsible for forming the spinal cord and paraxial mesoderm. Unlike NSC-based approaches, NMP-based protocols may increase the axial heterogeneity of in vitro models. This is due to one key intrinsic property of NMPs: their *HOX* gene expression code and progressive axial identity shift with extended culture time (Gouti et al. 2017). It has been shown that PSC-derived NMPs maintained in vitro for short periods of time (resembling early stages of development) primarily express anterior *HOX* genes, while NMPs kept in culture for longer, which resemble the transcriptomic profile of E9.5 mouse embryos, predominantly express posterior HOX genes (Gouti et al. 2017; Deschamps et al. 2017). It is thus likely that axial progenitor identity is acquired before neurogenesis begins and retained until the later stages of development, as reported in vivo (Metzis et al. 2018). In our study, we aimed to prime NMPs towards a neural lineage commitment, which is necessary for generating SCOs with high neural content. Recent NMP-based SCO protocols utilized for disease modelling result in a high percentage of mesodermal derivatives, as only a small subset of caudal progenitors develops into spMNs in these organoids (Faustino-Martins et al. 2020; Pereira et al. 2021). We successfully generated NMPs expressing *SOX2*, *TBXT* and *GBX2*, and demonstrated that a short and early RA pulse promotes their neural priming while maintaining their intrinsic caudal fate. Interestingly, we observed a reduction of SOX2 immunoreactivity in hiPSC-derived NMPs compared to NSCs, which fits previous data by Kelle and colleagues on the in vitro development of human axial stem cells (Kelle et al. 2024). Our findings are supported by in vitro differentiation of RA- primed mouse NMPs (Cunningham et al 2016), RA-primed human gastruloids (Hamazaki et al 2024) and by in vivo neurodevelopmental studies that showed that early exposure to RA during neural progenitor development controls NMP identity and maintenance (Kumar et Duester 2014, Cunningham et al. 2015, Durston et al. 1989, Blumberg et al. 1997). In summary, the effects observed in our study are likely due to RA acting as the signal that initiates the shift of NMPs into a "posterior" developmental mode, by activating the expression of 3’ *HOX* genes that govern posterior identity (Mahony et al. 2011, Nolte et al. 2019).Since the field standard protocol to differentiate mouse PSCs into spMNs was described two decades ago (Witcherle et al. 2002), numerous modifications of such model have been published and there is no consensus on how to best recapitulate spMN development in vitro. Some studies have compared the effect of different concentrations of key morphogens for spinal cord development on spMN patterning outcomes, such as WNT agonists, FGF2, RA and Smoothened agonists, (Andersen et al. 2020). Other reports have examined how the duration of caudalizing signals in vitro determines spMN axial identity (Lippmann et al. 2015; Mouilleau et al. 2021). Additional approaches focused on modulating the activity of the NOTCH signalling pathway in spMN progenitor maturation (Maury et al. 2015; Ben-Shushan et al. 2014; Tan et al. 2016). It is critical to consider which patterning molecules to use, at which concentration, timing and combination, as their corresponding signalling pathways directly influence the modulation of collinear *HOX* gene expression. Taking into account all the above evidence, we designed a protocol screen to systematically assess how patterning molecules commonly used in the field govern the generation of spMNs and the establishment of their specific axial identities. We began by comparing the original NSC-based approach with multiple variations of RA-primed NMP-based protocols. Importantly, to induce neuroectoderm fate on our hiPSC-derived RA-primed NMPs, we used only LDN during the majority of the GDF11 exposure phase to avoid blocking GDF11’s caudalizing effect with SB (Luo et al. 2019; Frohlich et al. 2022). In addition, and as developmental axial fate of CNS progenitors becomes relatively fixed after exposure to RA, we limited RA treatment window to 5-8 days and used it at a lower concentration than what is commonly used (50nM instead of 1 µM) to avoid stopping the HOX clock in the NPCs at anterior positions. Additionally, RA treatment of vertebrate embryos during gastrulation, when HOX gene expression is initiated, results in the rostralization of the CNS, skull defects and truncation of the axial skeleton (Conlon et Rossant 1992; Kessel et Gruss 1991; Marshall et al. 1992). We also tested including RA inhibitors intercalated with RA stimulation, as the specification of caudal HB9^+^ spMNs has been reported independent of RA exposure (Patani et al. 2011). Overall, while all novel protocol conditions generally resulted in increased caudal spMN fates compared to the field standard protocol, we did not observe major differences in spMN axial identity across them, particularly in terms of the order of delivery of neural induction, caudalization and progenitor expansion signals. This again highlights the intrinsic caudal bias of NMPs that does not seem to be heavily impacted by further exposure to morphogens. However, protocol condition #3 yielded a significant increase in the number of spMNs compared to the other paradigms. The early and prolonged exposure of neural progenitors to RA inhibitors characteristic of protocol #3 could explain this result, in agreement with a study reporting increased formation of OLIG2+ spMN progenitors and spMNs under similar conditions (Patani et al. 2011). Interestingly, in the mature SCOs, we observed a degree of regionalization within the axial spMN pools, with rostral and caudal spMNs spatially segregating in a self-organizing manner. This may be due to differential expression of surface proteins that provide migratory guidance to developing ventral spMN progenitors (Laussu et al. 2017).

Regardless of the patterning design, in our study, RA-primed NMP-derived SCOs generated only low numbers of rostral spMNs, which contrasts early reports on NMPs contributing to the formation of both paraxial mesoderm and the neural tube of all post-cranial axial levels in mice (Tzouanacou et al. 2009). Nevertheless, our results agree with more recent evidence supporting that after a certain time in culture, NMPs are unable to generate rostral neuroectoderm progenitors, even when treated with high concentrations of RA (Lippmann et al. 2015). Our data also agrees with studies showing that cervical neural tissues likely cannot be generated from neural-specific progenitors that originate from the axially-biased NMPs (Metzis et al. 2018, Nakamura et al. 2024, Bolondi et al. 2024). In addition, our NSC-derived SCOs, exposed to growth factors like GDF11 and FGF2 during their early development, contained few lumbar spMNs. According to literature, NSC exposure to RA at an early stage should completely halt the progression of the HOX clock towards caudal lineages and negate the effects of extrinsic caudalizing signals due to chromatin modifications (Lippmann et al. 2015; Mouilleau et al. 2021; Gribaudo et al. 2023). In differentiating mouse neural progenitors, RA activates RA receptors, which bind to rostral *HOX* chromatin domains, prompting a rapid removal of H3K27me3 and resulting in an irreversible cervical spinal identity (Mazzoni et al. 2013; Ahn et al. 2014). Based on our systematic analysis of numerous patterning paradigms showing that NSC-based SCOs still generate some caudal spMNs and NMP-based SCOs yet contain some rostral ones, we suggest that there may be some understudied overlap and plasticity between the processes of neurulation and axial identity acquisition, at least in vitro. We would like to emphasize, though, the importance of distinguishing between the intricate complexity of embryonic development and the inherent simplifications of cell culture models, which are designed to recapitulate specific developmental aspects. Discrepancies between in vitro and in vivo development -particularly when novel cell culture findings conflict with well-established developmental observations -are likely indicative of the need for further refinement of culture conditions rather than a misinterpretation of developmental processes.Once the generation of spMNs with a balanced RC identity was achieved, we aimed to enhance the physiological relevance of our CASCOs by incorporating endothelial-like cells and skeletal muscle. Our results showed that the presence of additional cell lineages seemed to provide crucial signals for promoting systemic neural maturation and the overall organoid health. Similar observations have been also reported by others, who demonstrated greater neural network activity and maturation in cerebral organoids transplanted in mouse cortex, likely through host-driven and engrafted endothelial vasculature (Mansour et al. 2018, Pham et al. 2018, Revah et al. 2022). No synergistic effect on the maturation parameters studied was detected when both cell types were added to the CASCOs versus only one of them. Worth noting, though, CASCOs containing both additional inducible cell lineages have ∼20% fewer cells available for neuronal commitment than the “CASCO+iETV2” or “CASCO+iMYOD” conditions. Similarly, the lack of synergistic effect on the overall organoid heath (ATP levels) may be also attributed to the significantly higher demands for ATP production of neuronal cells compared to other cell types (Leary et al. 1998, Du et al. 2008, Vinokurov et al. 2024).

Altogether, with this study, which leverages developmental cues that guide neural tube formation in vivo and compiles knowledge from the plethora of available protocols, we developed an advanced method for differentiating hiPSCs into physiologically relevant SCOs containing a wide axial variety of spMNs. We believe that this experimental model will be instrumental in studying the emergence and consolidation of human spMN diversity during spinal cord development, as well as whether and how certain motor neuron diseases might alter these processes.

## Acknowledgments

We would like to thank Thomas Becker (CRTD, TU Dresden) and Ana Rita Encarnação Aires (CRTD, TU Dresden) for their insightful feedback on our manuscript. We thank Lee Rubin (Harvard University) and Katrin Neumann (CRTD Stem cell Engineering facility, TU Dresden) for providing the MTAs enabling the work with the BJ-siPS WT, 1016A WT hiPSCs, 18A WT hiPSCs and the CRTD1 line, respectively, and the Sanger Institute for providing the KOLF-C1 hiPSC line. We thank Fabian Rost and all members of the DRESDEN-concept Genome Center (DcGC) (CMCB Technology Platform, TUD) for performing the scRNAseq experiment. We thank Silke White (Imaging, DZNE) and Ellen Geibelt (DZNE Imaging Platform and Light Microscopy Facility of CMCB) for their support and training on multiple microscopes and Zeynep Tansu Atasavum for training of our team on Arivis. We thank Anna Dalinskaya and Ines Rosignol for technical assistance. We thank Mike Karl (DZNE) for access to the Leica Cryostat and Michael Siewecke (CRTD) for access to the Applied Biosystem qPCR machine. We thank Gerd Kempermann (DZNE) for anti-NeuroD1 antibody and all past and present members of the Rodriguez-Muela Laboratory for their contributions to insightful discussions.

## Author Contributions

Felix Buchner: Conceptualization; data curation; formal analysis; visualization; methodology; writing, review and editing, funding acquisition. Zeynep Dokuzluoglu: conceptualization; data curation; formal analysis; review and editing. Joshua Thomas: data curation; formal analysis; methodology; review and editing. Antonio Caldarelli: data curation; formal analysis. Shrutika Kavali: formal analysis; methodology. Fabian Rost: data curation; formal analysis; methodology. Tobias grass: data curation; formal analysis, methodology; review and editing. Natalia Rodríguez-Muela: Conceptualization; supervision; writing, review and editing, funding acquisition; project administration.

## Methods and Materials

### Study approval and ethics

All experiments involving hiPSCs were performed in accordance with the ethical standards of the institutional and national research committees, as well as the 1964 Helsinki Declaration and its later amendments (General Assembly of the World Medical Association 2014). A positive evaluation by local ethics commission (Institutional Review Board) at the Technische Universität Dresden (Germany) was granted on 28th of Januaryy 2020 to use all the human iPSC lines (Reference Number: SR-EK 80022020).

### hiPSC culture

All hiPSCs lines (BJ-siPS, 1016A, KOLFC1, CRTD1, 18A) were thawed in presence of 10µM ROCKi, grown on P6 dishes (Corning), which were coated 40 min at RT with Matrigel-DMEM (Corning, Gibco). Cells were cultured in mTeSR plus (STEMCELL Technologies), which was refreshed every 2-3 days. Splitting of the hiPSCs was performed with 1 min incubation in 0.5M EDTA, once 60- 80% confluency or terminal size of the colonies was achieved, then cells were washed with 1xDPBS, scraped off the plates and passaged in clumps. Cell lines were used for differentiation into spMNs, karyotyped and sequenced before (Boulting et al. 2011, Ng et al. 2015, Rodriguez-Muela et al. 2017, Rodriguez-Muela et al. 2018, Hildebrandt et al. 2019, Voelkner et al. 2022). Healthy hiPSCs were processed in clump-based dissociation, centrifuged for 5min at 1,000rpm and then transferred to CryoStor® CS10 (Stemcell), in Cryovials (Sigma Aldrich), and to a Mr. Frosty™ Freezing Container (Nalgene) and ultimately stored at -80°C for 1-2 days. Afterwards, Cryovials containing iPSCs were stored in liquid N2 tanks long term.

### CRISPR-Cas9 mediated tagging of the endogenous ISL1 locus

To generate Islet1 reporter line we followed a two-vector based knock-in approach. Briefly, one plasmid (px458, Addgene Plasmid #48138) expressing sgRNA as well as Cas9 was used to generate pTG-Cr- ISL1 and to introduce double-strand breaks near the stop codon of Exon 6 of Islet1 locus. To do so, a CRISPR guide (gRNA) with an estimated cleavage site right next to the stop codon of exon 6 of the *ISL1* locus were designed and single-stranded (ss) oligos (IDTDna) (listed in the Materials and Methods Table) were annealed and cloned into the px458 plasmid (Addgene # 48138) using BbsI and T7 DNA ligase in a one-step digestion-ligation reaction. Correct insertion of the ssOligos into px458 plasmid was confirmed via Sanger Sequencing (Microsynths). ssOligo Info see extended methods table and gBlock sequence available upon request. The second vector pTG-HR-ISL1-P2A-mNeonGreen was derived through modifications of the standard HR120-PA-1 vector (Systembio). To create a new multicistronic vector the copGFP- polyA cassette was removed via digest using EcoRI and NruI and the vector sequence was restored via gBlock cloning which introduced a P2A cassette. In subsequent steps the XhoI site and Gibson assembly was used to introduce a gBlock consisting of mNeonGreen CDS followed by a WPRE element. For introduction of the 5-prime homology arm the plasmid was digested with NHEI and a gBlock (IDT) consisting of a 429-bp sequence, encoding part of intron 5 as well as the last exon of the *ISL1* gene excluding the stop codon was cloned into the vector via Gibson assembly (NEB) (and in frame with p2A- mNeonGreen sequence followed by the WPRE site, loxP-flanked drug-resistance cassette and fluorescent marker mRuby). Next, the vector was digested with BamHI and the 3-prime homology arm containing of 421-bp homologous to intron 6 of the *ISL1* locus was cloned into the vector via Gibson assembly, resulting in pTG-HR-ISL1-p2A-mNeonGreen. For the generation the ISL1 reporter line cells were nucleofected with pTG-Cr-ISL1 and pTG-HR-ISL1-p2A-mNeonGreen plasmids using a 4-D nucleofector system (AMAXA) and the P3 Primary Cell 4D-Nucleofector Kit (Lonza) following manufacturer’s instructions. 2 days later succesful targeted cells were selected with puromycin for a total of 7 days, followed by nucleofection with pCAG-Cre:GFP plasmid (Matsuda and Cepko, 2007) to remove the loxP-flanked drug-resistance cassette and flurescent marker mRuby. 4 days later cells were FAC-sorted and mNeonGreen^+^/mRuby- cell population was enriched and plated at clonal densities on freshly Matrigel-coated cell culture dishes. 24 individual colonies were picked, expanded, gDNA was extracted using DirectPCR lysis reagent (CELL, Viagen) and Proteinase K (Viagen) and succesful targeting was confirmed via PCR strategy using primer strategy recently described by us (Grass et al, 2024). Out of the succesful targeted clones (data not shown), 2 were expanded and used for experiments. Validation of ISL1-p2A-mNeonGreen reporter was proven via co-immunstaining with commercially available antibody (Abcam), resulting in more than 90% overlap.

### Transposon-mediated gene editing of WT BJ-sIPS hiPSCs

Plasmids were propagated using One-Shot TOP10 Bacteria (E.coli) or VB Ultra Stable Bacteria (E.coli) with LB medium. Firstly, the BJ:ETV2i- hiPSC line was generated by co-transfecting WT BJ siPS with 4 μg of pBAN-Puro-TT-hETV2-isoform2 (ENST00000402764.6) and 1 μg of pRP[Exp]-mCherry-CAG>hyPBase vectors, similarly as described elsewhere (Ng et al. 2021). Selection was performed using 1 μg/ml puromycin for two weeks prior to clonal selection. Briefly, and as described before by others, the BJ:Baf60c-MYOD1-hiPSC line was generated by co-transfecting WT BJ siPS with 2.5 μg each of the ePB-Bsd-TT-BAF60c and ePB-Puro-TT-m-MyoD vectors, along with 1 μg of pRP[Exp]-mCherry-CAG>hyPBase (Lenzi et al. 2016, Tiago et al. 2021). Selection was performed using 1 μg/ml puromycin and 5 μg/ml blasticidin for two weeks and gave rise to stable and doxycycline inducible cell lines. Afterwards, we performed single cell cloning as described elsewhere (Grass et al. 2024).

### hiPSC aggregate formation and SCO patterning

Once hiPSC colonies reached terminal size and plates showed about 60-80% confluency, they were processed for further experiments. For aggregate generation, hiPSC colonies were first washed with 1xDPBS and treated 1 min with ReLeSR solution at room temperature and then lifted off the plates, using Corning® Cell Lifter 3008 (Corning). Cells were immediately transferred into 5ml of mTesR (StemCell) medium and dissociated into clumps. *<u>For SCO</u> <u>protocol screen</u>*. The cells were transferred into in P10 ultra-low attachment Petri dishes (Corning), containing 10ml of mTesR medium, ROCK inhibitor Y-27632 (StemCell) and 20ng/ml recombinant FGF2 (Qkine). After 24h, half of mTesR medium was refreshed and 20ng/ml FGF2 was added. EBs were grown for another 24h. EBs were continuously shaken at 60rpm inside of an incubator (37°C, 5% CO2). After three days in culture, the EBs were transferred into mTesR containing the cocktail CEPT (Chen et al. 2021, Ryu et al. 2023), and shaken at 60 rpm for another 18h. The EBs were then transferred into NIM medium containing the WNT agonist CHIRON99021 (Tocris) at 3μM and 20ng/ml FGF2. After 48h, 25nM of RA (Sigma-Aldrich) was added for 2h. Lastly, NIM was replaced and NMP-like aggregates were cultured in NIM with 20ng/ml FGF2 for 12h. They were then transferred onto P24 ULA Ultra Low attachment plates (Corning) to start the patterning protocols (or drug treatments). *<u>For CASCO formation</u>*. Instead of clump formation, adherently patterned cells were treated with Accutase (Corning) for 5min at 37°C, viable cells were counted with a Neubauer Chamber and 10,000 cells of adherently patterned NSCs and NMPs were mixed together at a ratio of 1:1 in P96 ULA plates (Corning) and spun down at 120g for 3 minutes. These were left in NIM+CEPT+FGF2+SAG for one day to recover before neural induction with SB+LDN in NIM, and then successive patterning steps continued. Concentration of small molecules for SCO protocols used: 10 μM SB431542 (Stemgent), 1μM LDN193189 (Stemgent), 3μM CHIRON99021 (Tocris), 1μM RXR inhibitor UVI 3003 (Tocris), 1μM RAR inhibitor AGN 193109 (Tocris), 1μM, 50nM or 25nM RA (Sigma-Aldrich), 200ng/ml FGF8 (Peprotech), 10μM Notch-inhibitor DAPT (Tocris), 120ng/ml FGF2 (Qkine), 1μM Smoothened Agonist SAG (Merck), 50ng/ml GDF11 (Peprotech), 10ng/ml BDNF (Qkine), 10ng/ml GDNF (Qkine). Neural Induction Media (NIM) for SCO culture contained 477.5 ml Advanced DMEM and 477.5 ml Neurobasal Media (both from Life Technologies), 10 ml Penicillin-Streptamycin and 10 ml B27 supplement with Vitamin A (Thermo Scientific), 10 ml GlutaMax (Gibco), 1.82 ml ß-Mercaptoethanol (Sigma-Aldrich), 5 ml N2 supplement (Thermo Scientific), 8 ml 20% glucose in 1xDPBS (sterile filtered from Sigma Aldrich), 200 μl 5,000x Ascorbic Acid (StemCell), and 1 ml 1,000x Antioxidant Supplement (Thermo Scientific).

### High content live imaging

Live imaging of SCOs derived from healthy reporter line BJ- ISL1:mNeonGreen under different patterning approaches was performed with an Operetta CLS (PerkinElmer, USA). Images were taken every 6 hours for multiple days, using a 10x Air Objective. NIM with small molecules was briefly exchanged for Neurobasal Medium without Phenol Red (Thermo) and after imaging, the respective patterning medium was supplied again. SCOs were counterstained for 90min with 5µM Calcein Red dye at 37°C and re-stained at day 6 and day 12 of the differentiations. Mean fluorescence intensity of ISL1:mNeonGreen was automatically quantified with high-content imaging analysis software HARMONY (PerkinElmer), restricted to the EBs identified in Brightfield (BF) or by CellTrace™ Calcein Red-Orange (Invitrogen) dye.

### RNA isolation, cDNA synthesis, primer Design and qRT-PCR

After two successive washes in 1xDPBS, total RNA and protein were extracted from the RNA/Protein NucleoSpin Kit (Macherey Nagel) or TRI Reagent (Zymogen), following the manufacturer’s instructions. The following adjustments for the Nucleospin extraction were meade: In the lysis buffer RP1, 1xHalt-Protease Inhibitor (ThermoScientific) and 2x ß-Mercaptoethanol (Sigma-Aldrich) were added, with subsequent RNA purification involving four instead of two washes in ice-cold RA3 wash buffer and incubation of RNA in 70% ethanol with 20µl of 3M Sodium Acetate (pH adjusted to 5.2 with Glacial Acetic Acid) for 5 minutes. Double elution with 20- 30µl of Nuclease-free Water (Promega) was performed. The following adjustment for the TRI Reagent extraction were made: Glyogen (Thermo Scientific) was added to the aqueous phase after Chloroform-mediated separation of the Phenol at 1µg/µl to enhance yield. RNA concentration and quality was measured using NanoDrop TM 1000 Spectrophotometer (Thermo Scientific). For cDNA synthesis, 250- 500ng of RNA was reverse transcribed using the Applied Biosystems™ High-Capacity cDNA Reverse Transcription Kit in a C1000 Touch Thermal Cycler (Biorad), following the manufacturer’s instructions. Primers for quantitative reverse transcription PCR (qRT-PCR) were designed using PrimerBLAST NCBI (Ye et al., 2012) and controlled for dimer formation using IDT OligoAnalyze (IDT) and validated using pooled cDNA from hiPSCs and SCOs at different developmental stages. qRT-PCR was conducted using the GoTaq qPCR Master Mix (Promega) on a 384-well QuantStudio™ 5 Real-Time PCR System (Applied Biosystems) in Comparative Cт (ΔΔCт) mode with CXR reference dye, analyzing gene expression levels relative to the housekeeping gene h18S. Data was analyzed with the ΔCq method according to Pfaffl (2001) and ultimately in PRISM (Graphpad) or R Studio (R). Gene expression is indicated as LOG2FC of ΔCq or FC with respect to the housekeeping gene h18S.

### Dissociation and scRNAseq of NSC-derived SCOs

WT BJ hiPSC aggregates were patterned as described in Fig1A and at day 20 they were dissociated as described in Grass et al 2024, using a papain/DNase solution (Worthington, LK003178, LK003172). Cell viability was determined using trypan blue staining and 50K cells per sample were pooled at a final concentration of 2K cells/µL in 0.04% PBS-BSA. Droplet-based scRNA-seq was performed using the 10x Genomics Chromium Single Cell Kit v3. Raw sequencing data was processed using the count command of Cell Ranger (v7.0.0) provided by 10xGenomics. To build the reference, human genome GRch38/hg38 and gene annotation Ensembl104 were downloaded from Ensembl. The annotation file was filtered using mkgtf command of Cell Ranger (options:attribute=gene_biotype:protein_coding,attribute=gene_biotype:lincRNA,attribute=gene_biotyp e:antisense) then using the filtered annotation and genome sequence as input to the mkref. command to build the reference. FASTQ files were processed with the count command, which aligned reads to the reference, generating a gene-barcode-matrix, representing gene counts per cell. For further bioinformatical analysis, the dataset was filtered to exclude low-quality cells (criteria: nFeatures > 1000 and percent of mitochondrial genes < 10%). We analyzed the gene expression profiles of 12,637 cells. On average, we mapped 95.6% of all reads to the human genome and determined 586,951,087 unique reads, accounting for 5,067 median genes per cell. Next, we applied normalization, scaling, and dimensionality reduction using standard Seurat functions (Sajita et al 2015). For normalization, gene expression values were adjusted by total UMI counts per cell, multiplied by 10,000 (TP10K), and then log-transformed using log10(TP10K + 1). Dimensionality reduction was performed using principal component analysis (PCA) on the top 2,000 variable genes identified via the vst method. For two-dimensional data visualization, uniform manifold approximation and projection (UMAP) was computed using the first 20 principal components (PCs). Cell clustering was carried out using the Louvain algorithm, also based on the first 20 PCs, with a resolution set at 0.1, generating Dotplots to show LOG2FC of gene expression per cluster. Marker genes specific to each cluster were identified with the Wilcoxon rank sum test, utilizing the FindAllMarkers Seurat function (parameters: only.pos = TRUE, min.pct = 0.20, logFC.threshold = 0.20, min.diff.pct = 0.1). The biological identity of each cluster was inspired by previous cluster annotations from hiPSC-derived SCOs (Grass et al., 2024) and from published studies on spinal cord gene expression (Stifani et al. 2014; Sagner et al. 2019, Andersen et al 2023, Catela et al., 2024). The confusion matrix was calculated to represent the frequency of cells from each group within each cluster. All visualizations were created using standard Seurat functions along with custom ggplot2 (v. 3.4.0) scripts. The code utilized in this study will be made available on GitHub upon manuscript acceptance. Seurat objects and processed data files will be available upon request.

### Cryosectioning, immunofluorescence staining, imaging and image processing

After o.n. fixation of organoids in 4%PFA in 1xDPBS, organoids were incubated for 15 min at RT in 0.5M Glycine-PBS solution to quench aldehyde groups. Then, they were subjected to cryoprotection in 15%Saccharose/PBS for 48h and 30% Saccharose/PBS, flash-frozen on dry-ice in TissueTek OCT embedding compound (Sigma) and stored at -80°C. Cryoblocks were sectioned at 15µm thickness with the Cryostat Leica CM3050S (Leica) and sections collected onto Thermo Scientific™ SuperFrost Plus Slides (Thermo). Slides were dried at 37°C on a TFP40 Stretching Table (MEDlite) for 10min before extended storage at −80°C. For immunofluorescence staining of organoid sections, slides were thawed at RT for 5 min, rehydrated in 1xDPBS for 10min and post-fixed in 4% PFA (ChemCruz) in 1xDPBS for 10 min. Slides were washed in 1xDPBs twice and blocked for 1 h with Blocking Buffer (BB) (0.25% TritonX, 4% NGS or 3% BSA, 0.1% Tween20 in 1xDPBS) at RT in a humid chamber. Afterwards, sections were circled with an Elite Pap-Pen (Diagnostic BioSystems), slides were washed once for 10min in DPBS and Primary Antibodies in BB were added for 2h at RT. Slides were washed in 1xDPBs twice (10min each) and Hoechst33342 nuclear dye was added at 2µg/ml for 10min. Slides were washed in 1xDPBs twice, mounted with Vectashield Antifade Mounting Medium (VectorLabs) on HighPrecision (170±5µm) coverslips (Carl Roth) and sealed with clear nailpolish. All washing steps in 1xDPBS were carried out at RT on a neoLabLine Multi Shaker (neoLab) at 100rpm. Imaging was carried out with a Zeiss Confocal Spinning Disc, using automatic scan of manually selected organoids/tile regions, with a 20x Air Objective unless indicated otherwise. Z-Stack scans were recorded for each organoid section, Niquist theorem was always considered, to select optimal interval size in z-direction. For imaging of the organoids in the protocol-screen, unbiased-automated slide scanning was performed on a Zeiss Axioscan.Z1 microscope, using a 20x Zeiss Plan-Apochromat Air objective. Extended Depth of Focus option (EDF) was used for recording Z-Stack scans of at least 12 µM thickness with an interval of 1 µm. Maximum Intensity Projections were made and tiles stitched with 15% tile section overlap by Hoechst33342 as reference channel. All Images were sharpened by 20%, adjusted for contrast, brightness and gamma, using always the same LUT setting an intensity cutoffs for background determination for each group. Automatic quantification of SCO sections was carried out in Arivis Vision4D 3.3.0 (Zeiss) after batch-transcribing the files with ArivisSIS converter (Zeiss). Briefly, CZI.files were converted into arivisSIS format with no compression. The quantification pipelines in Arivis Vision4D 3.3.0 usually included: 1) A percentage-based Threshold for the channel of interest, restricting histogram values. 2) Global Enhancement “Simple Sharpening Filter" or Richardsson-Lucy-Deconvolution. 3) Usage of BlobFinder- Nuclear Segmentation Tool for identifying Nuclear Shapes with Diameter >12µM, Probability threshold=4%, Split sensitivity=80%, slightly adjusted dependent on the marker of interest 4) Pixel-based Compartment Analysis with “Fill-Holes”-option, segmentation in x,y,z-direction to identify cells that are double positive for certain HOX markers and ISL1(2). 5) Segment operation “Feature- Filter" on Surface-Area (µm²) > 60 µm², VOXEL > 150 µm3, Sphericity > 0.6. HOXC10 protein can be present both in nucleus and cytoplasm (Pathiraja et al 2014). In our analysis HOXC10 signal was assessed in volumetric overlap with nucleus staining (Hoechst). We proceeded similarly regarding other HOXs, always setting at least 70-80% nuclear volumetric overlap as an analysis parameter. For analyzing MAP2 staining and Nestin staining, we used Frangi’s vesselness measure (Frangi et al. 1998). For PSD95/MAGUK, BDNF and CHAT we trained a machine-learning model based on Illastik for minimum 150 features and hosted the model in Arivis (Berg et al. 2019). Quantification was carried out in an unbiased way, always using “Batch Analysis" option for all organoids. Dispersion around the mean or median is shown always either as SEM or interquartile range, respectively. Analysis outputs were exported as CSV files, and then compiled using a custom R-code, available on request. Final analysis values were sorted using Microsoft Excel 2016, normalizing the Hoechst33342 cell count number for each organoid, performing one-point normalization to the average relative value for the control group, and plotted using Graphpad PRISM. The distance map was created in Zen by drawing a peripheral line, using the “Profile’’ option, showing separation of post and presynaptic marker staining. All Images were assembled in Affinity Designer 2.

### CellTiter Glo Assay for ATP measurement in SCOs

The ATP assay was conducted following the manufacturer’s instructions using white opaque walled microwell assay plates containing 1-3 SCOs per condition. CellTiter-Glo® Reagent was added in an equal volume to all wells, followed by mixing the contents for 2 minutes on an orbital shaker to induce cell lysis. Finally, luminescence was recorded using an Infinite 200 Pro Reader. Signal in RLU was normalized against an ATP-Standard curve (1μM to 10nM), with freshly prepared dilutions every time.

### Statistical analysis of measurements

We routinely carried out Shapiro-Wilk test for normality assessment of data points. Only if all experimental groups were Gaussian distributed, we proceed with parametric tests (Starbuck et al 2023). Outlier detection was set with the lower limit at 2 standard deviations below the mean (μ - 2σ) and the upper limit at 2 standard deviations above the mean (μ + 2σ). To analyse median changes of LOG2FC of gene expression, we regularly used a one-tailed Mann-Whitney U test (Nachar 2008, Mann et Whitney 1947) due to the non-normal distribution in some qPCR experiments. For experiments with higher number of data points and Gaussian distribution of the data, we conducted a one-way ANOVA followed by Fisher’s LSD post hoc tests for pairwise comparisons (Stuhle et Wold 1989, Meier 2006). If needed, data were analyzed using a two-way Mixed Model (Ato García et al 2013) or two-way ANOVA with Fisher’s LSD Test, respectively. For the analysis of related cell counts during assessment of the organoid protocol screen, we used the non-parametric Kruskal-Wallis test (Kruskal et Wallis 1952) and uncorrected Dunn’s test (Dunn 1961). Regarding the analysis of cell numbers in mixed organoids in Fig. 4-5, given the complexity involving multiple experimental conditions and various cell lines, we opted for a two-way ANOVA with Tukey-Kramer post hoc test (Yates 1934, Fujikoshi 1993) to facilitate reliable pairwise comparisons between experimental conditions with unequal sample sizes and variant numbers of samples (Driscoll 1996). To assess the difference in synaptic puncta (Synaptophysin, Pan-MAGUK) and ChAT/BDNF signal, we conducted Unadjusted, one-sided and pairwise t-Test with Welch correction for multiple comparisons due to unequal variances between conditions (Welch 1951). Exact information about biological and technical replicate number is indicated in each figure legend.

## 14. Methods - Reagents and Tools Table

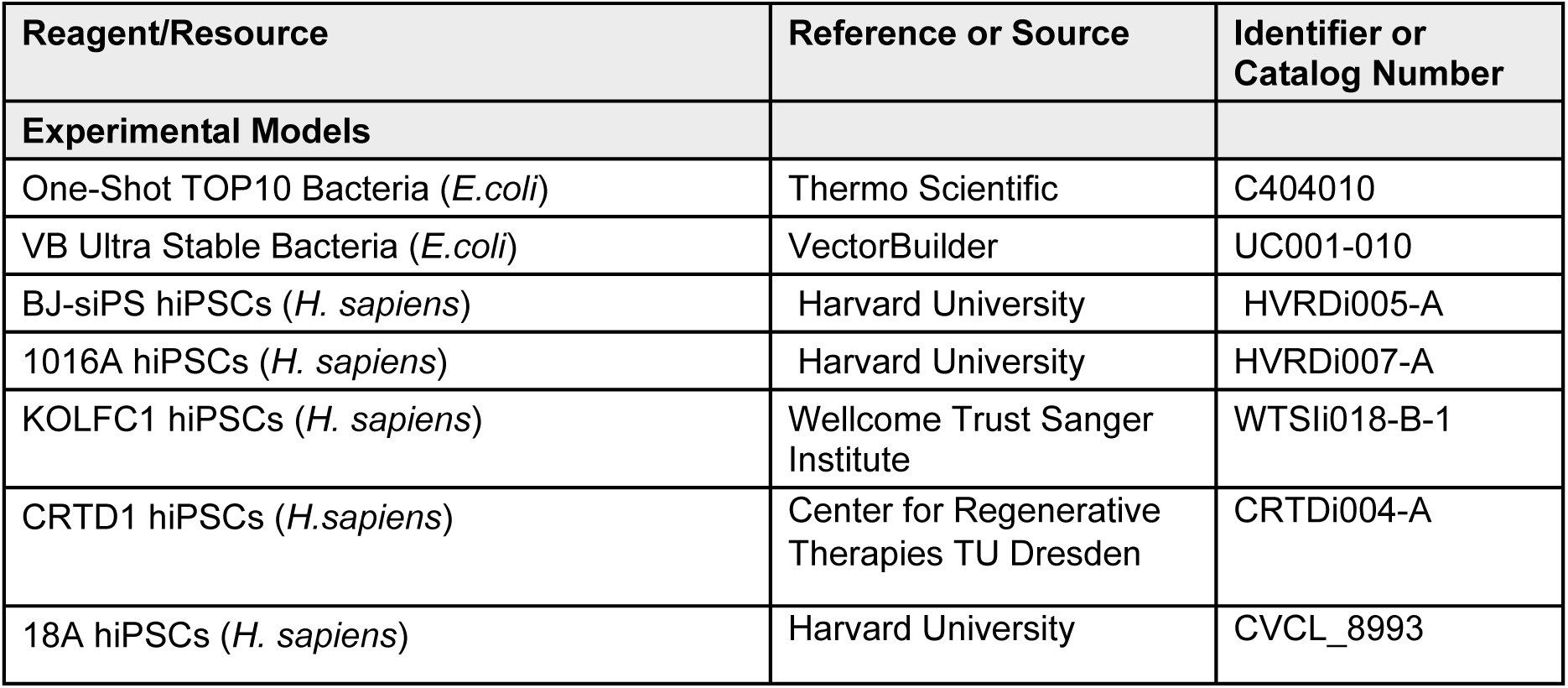
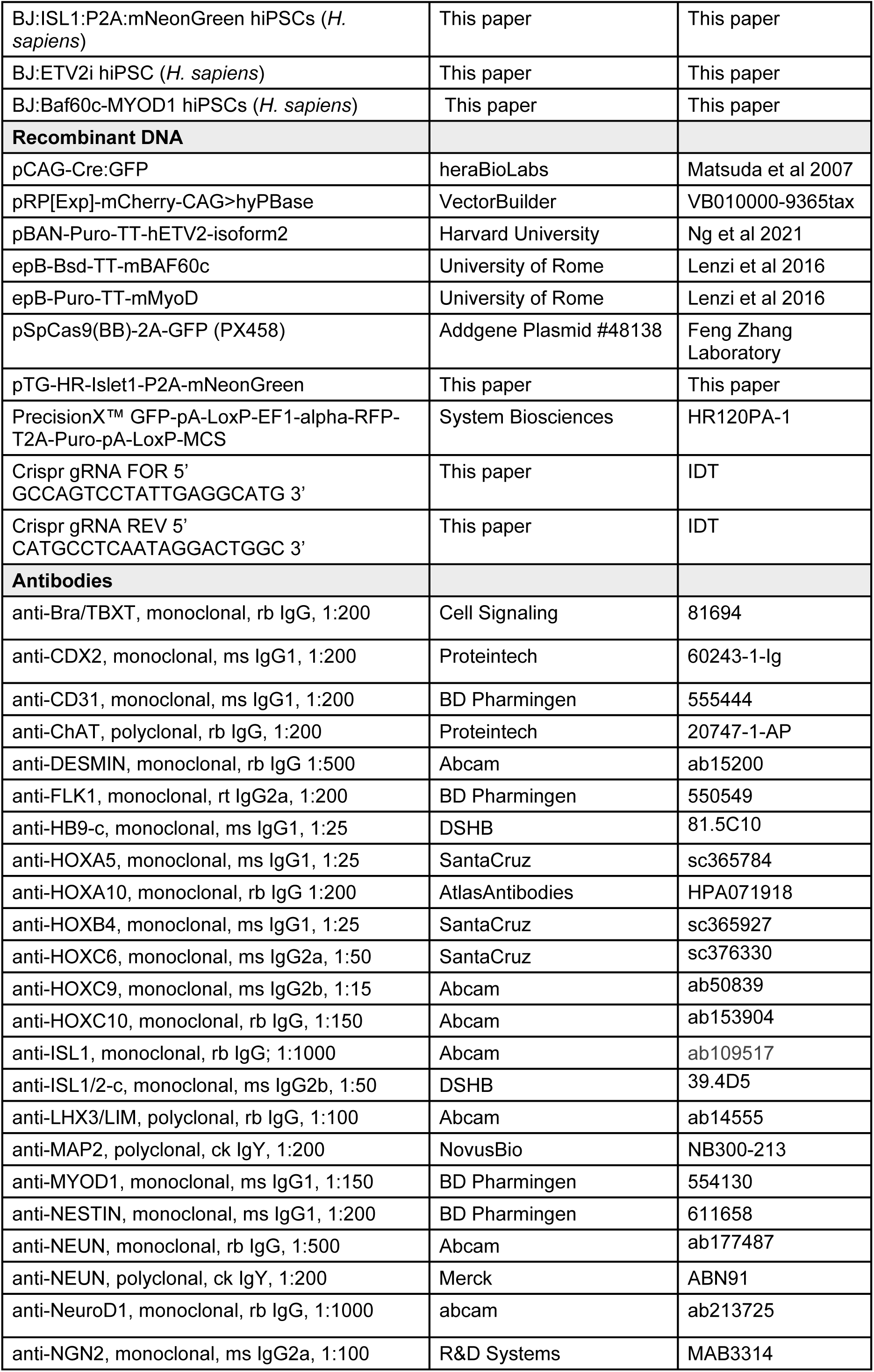
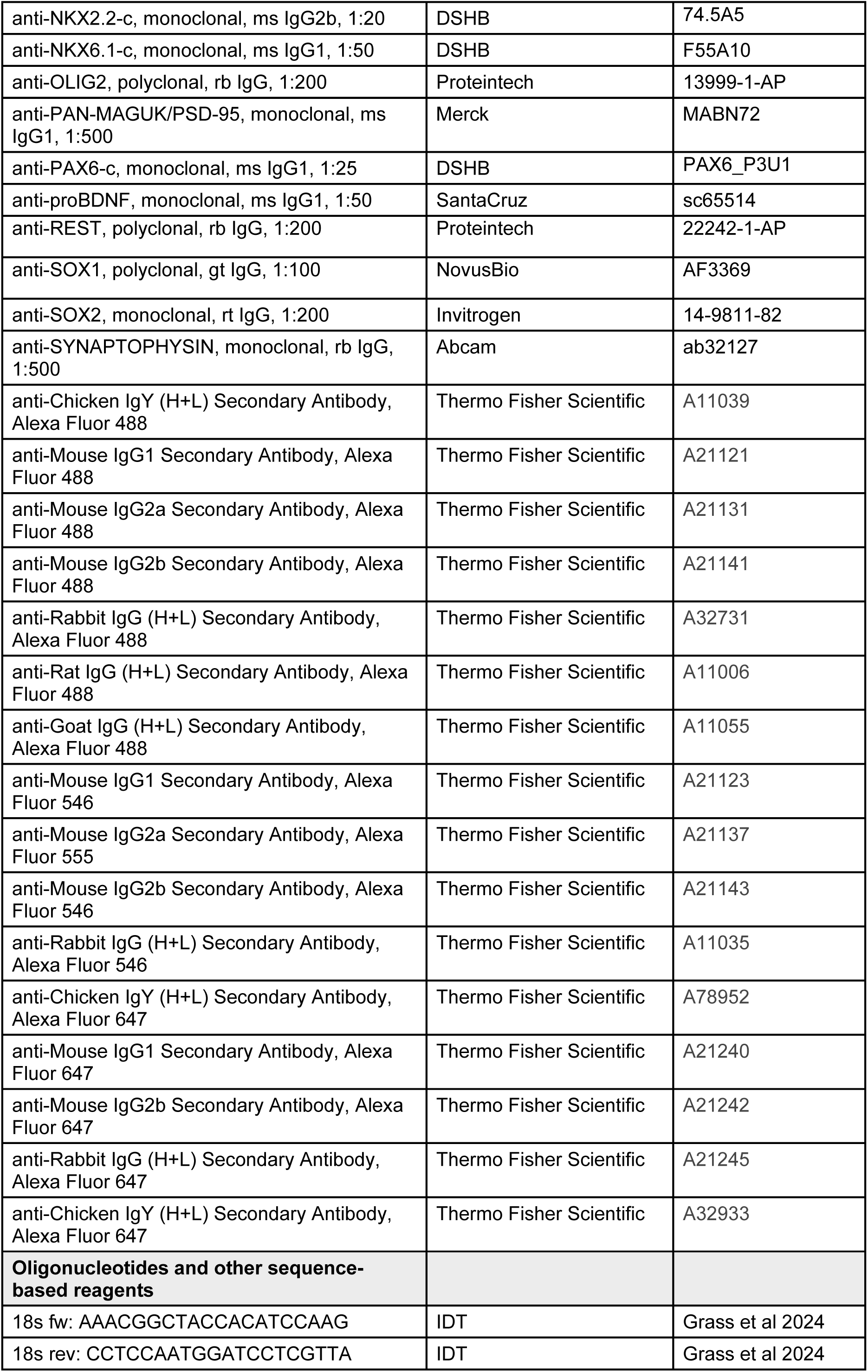
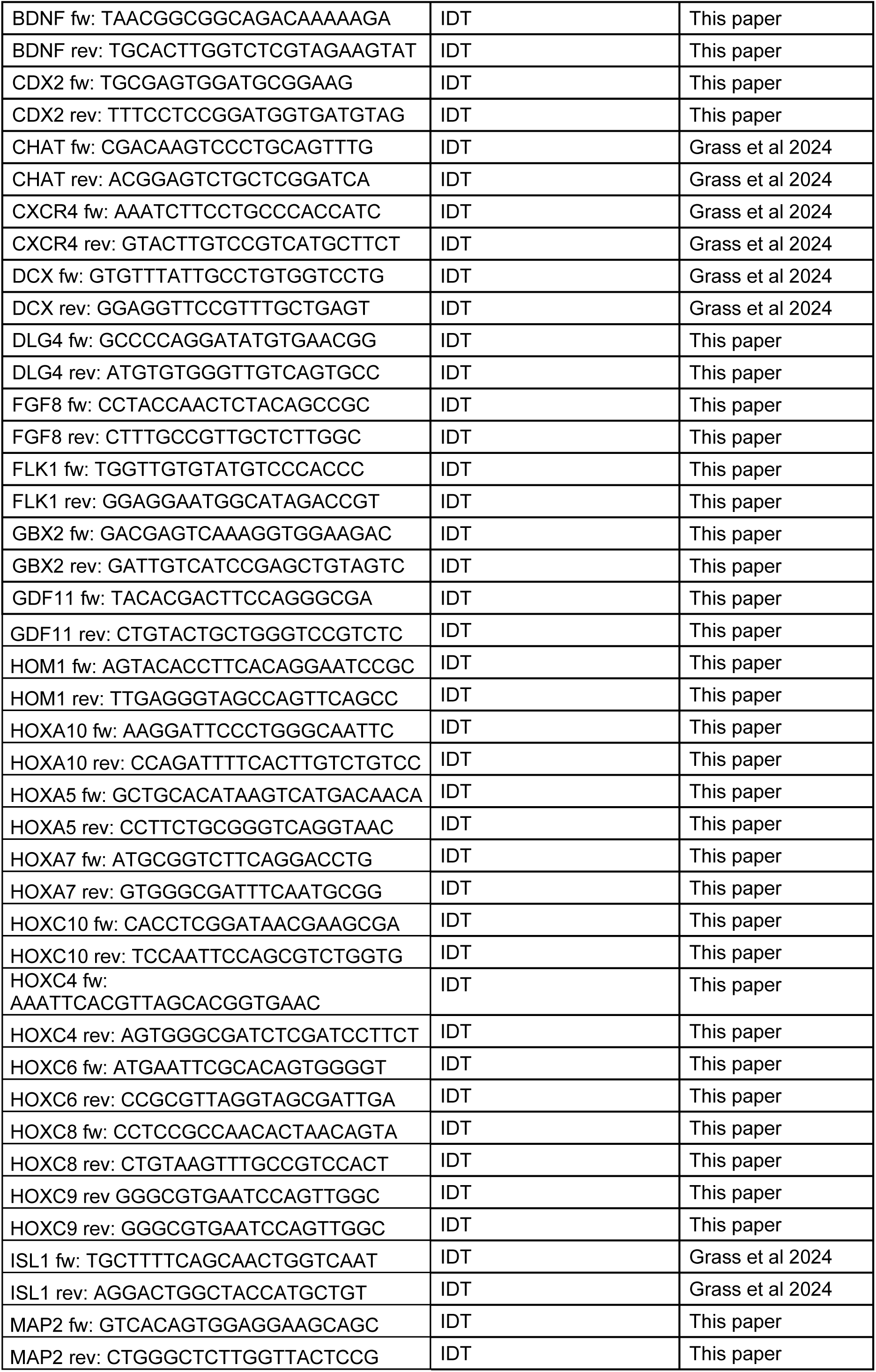
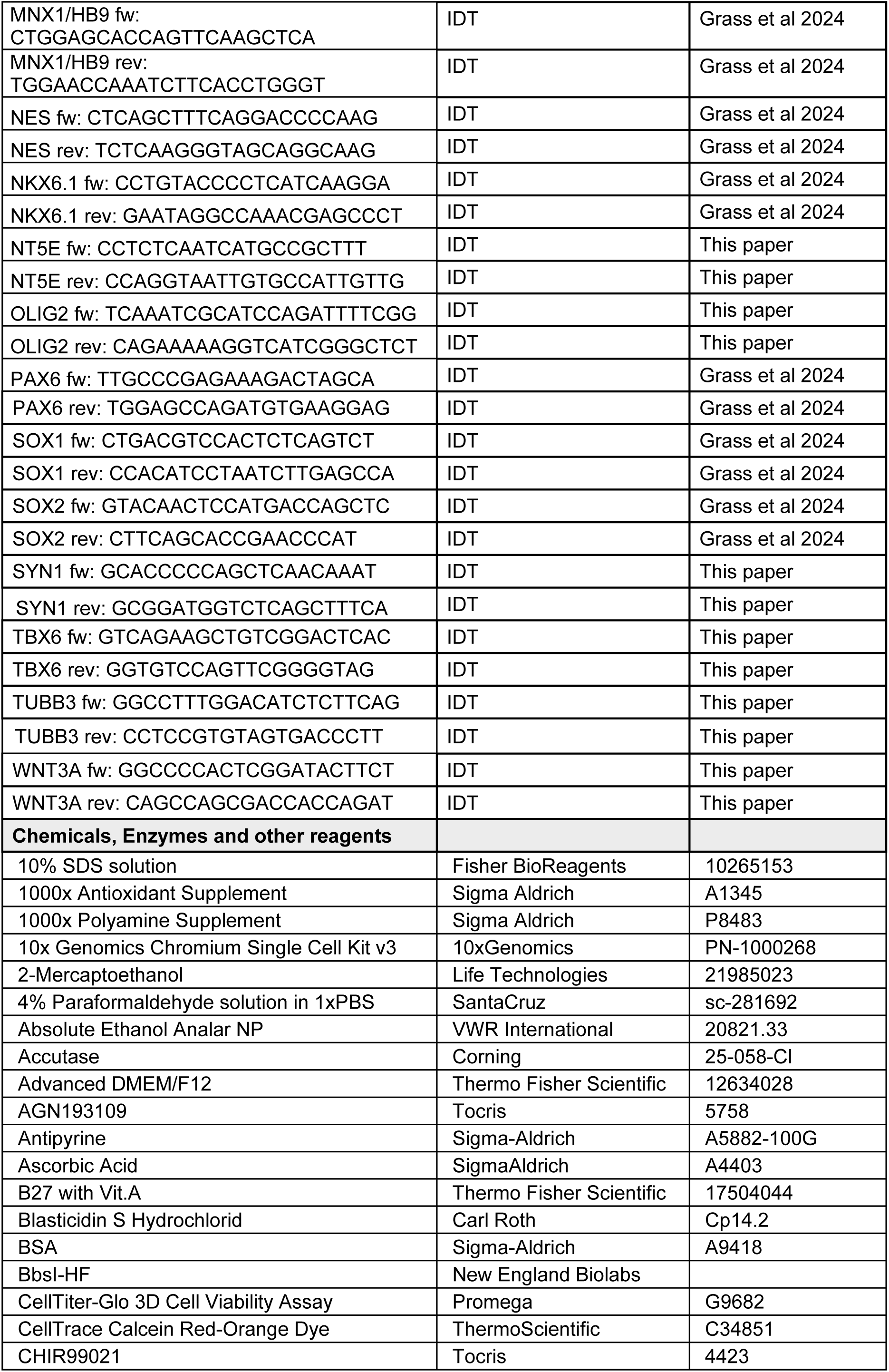
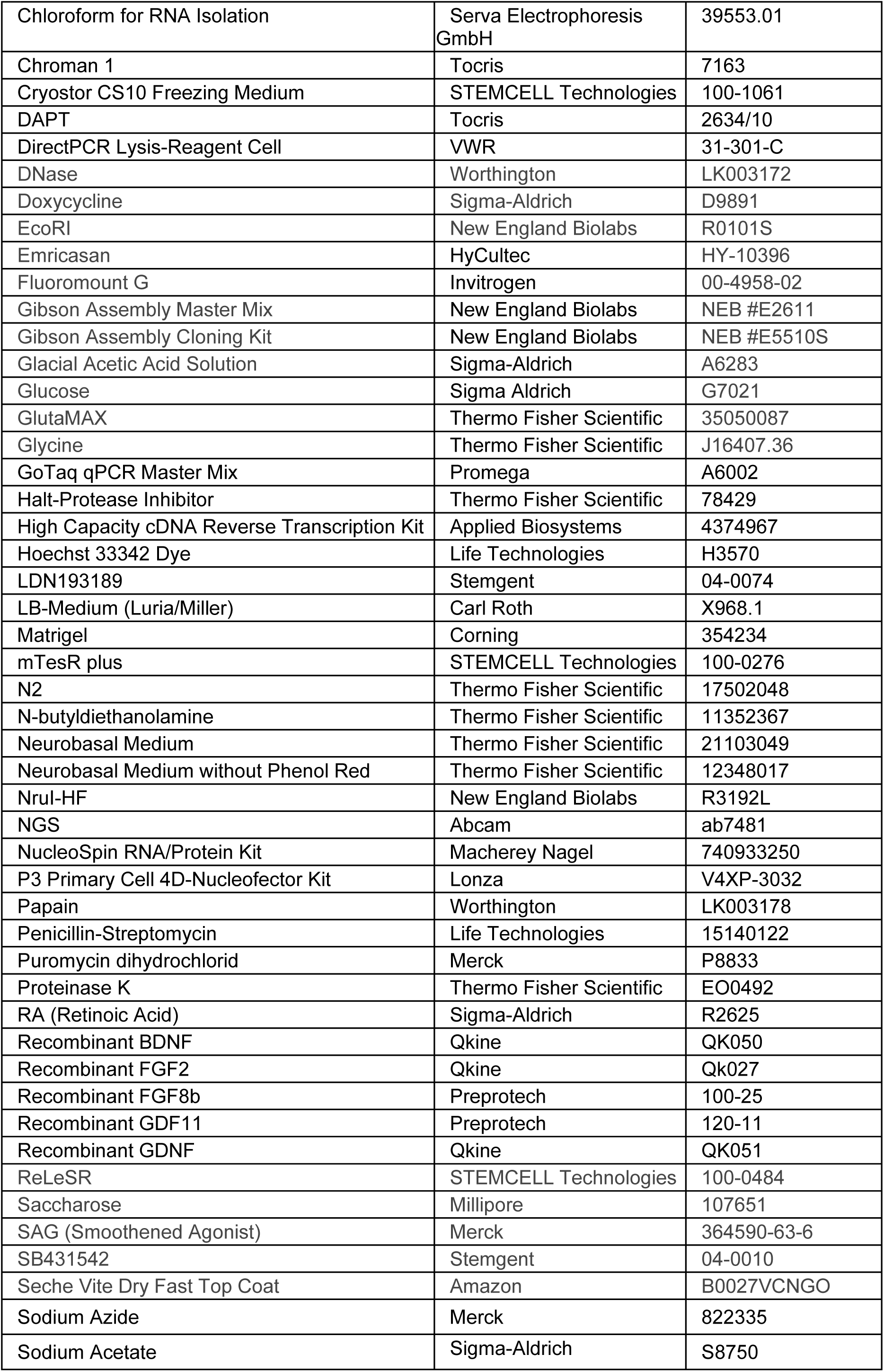
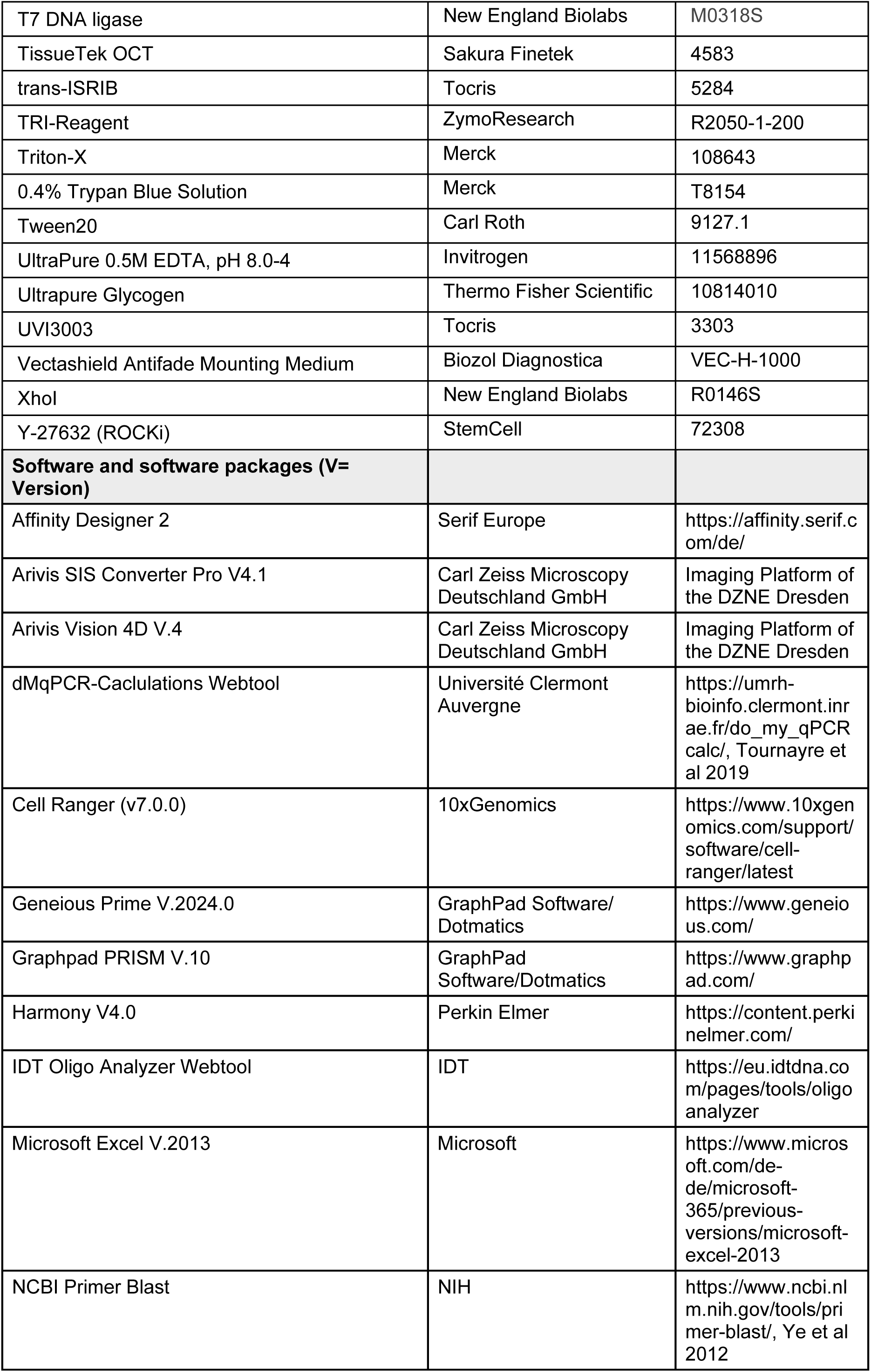
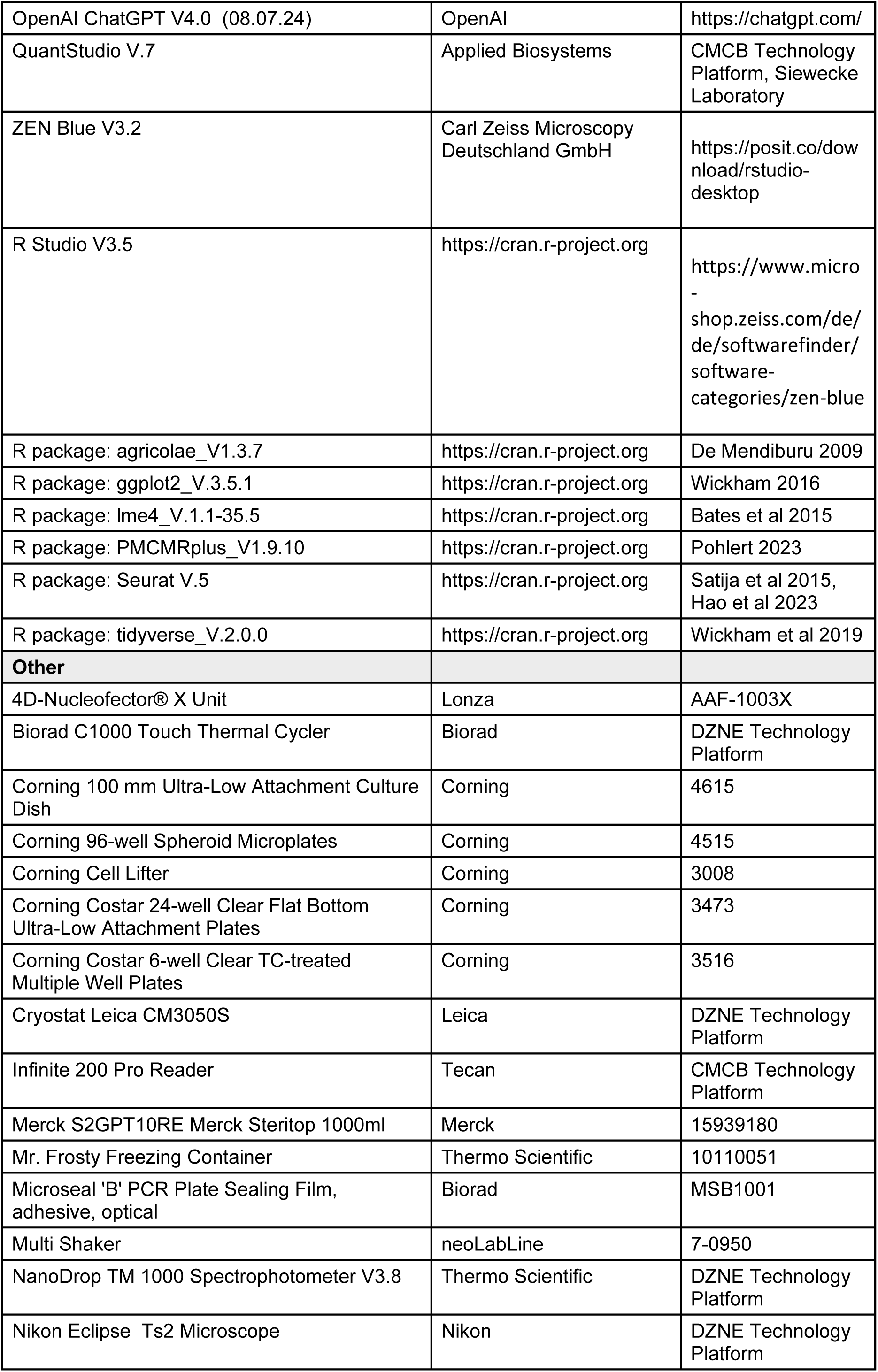
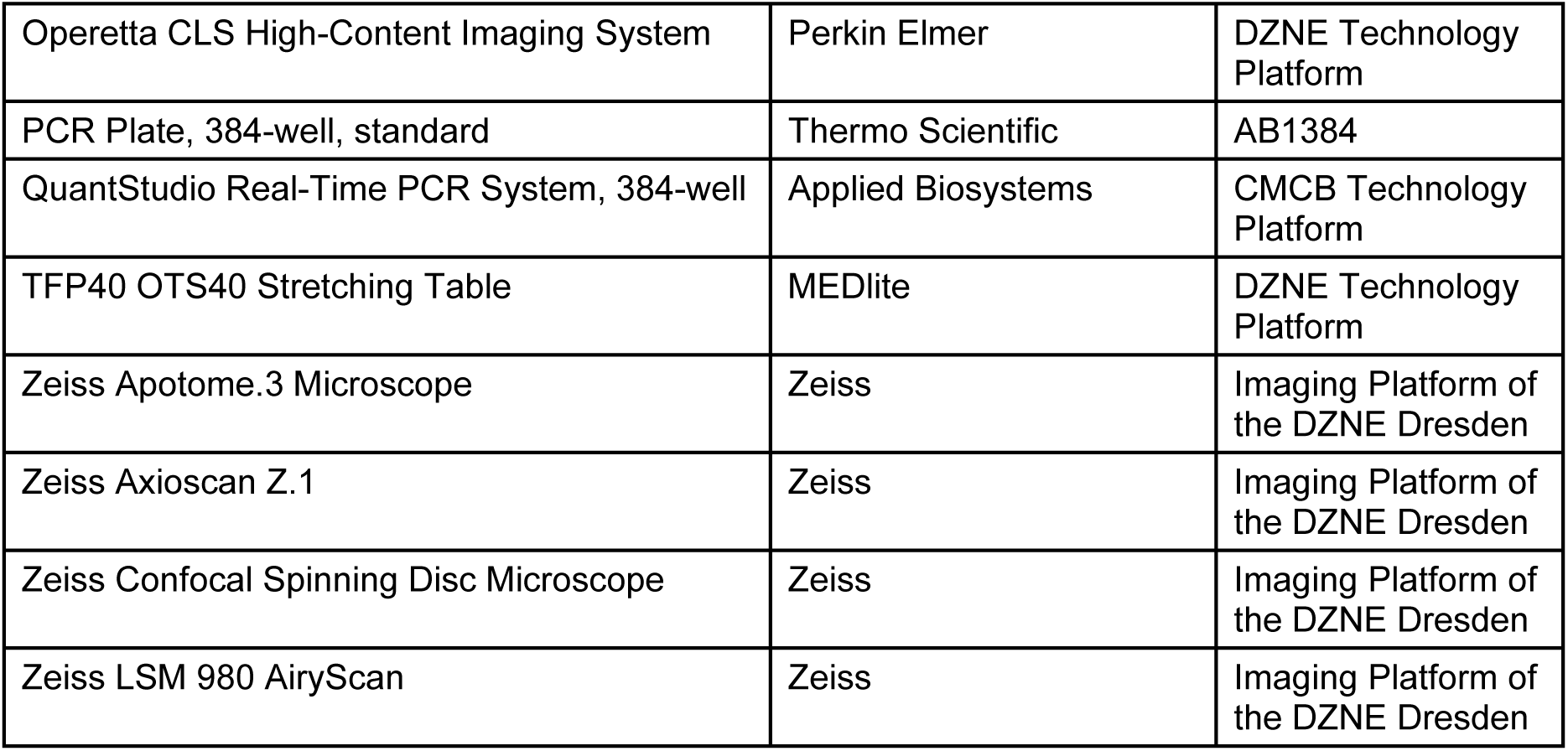

---

**Supplementary Figure S1.**
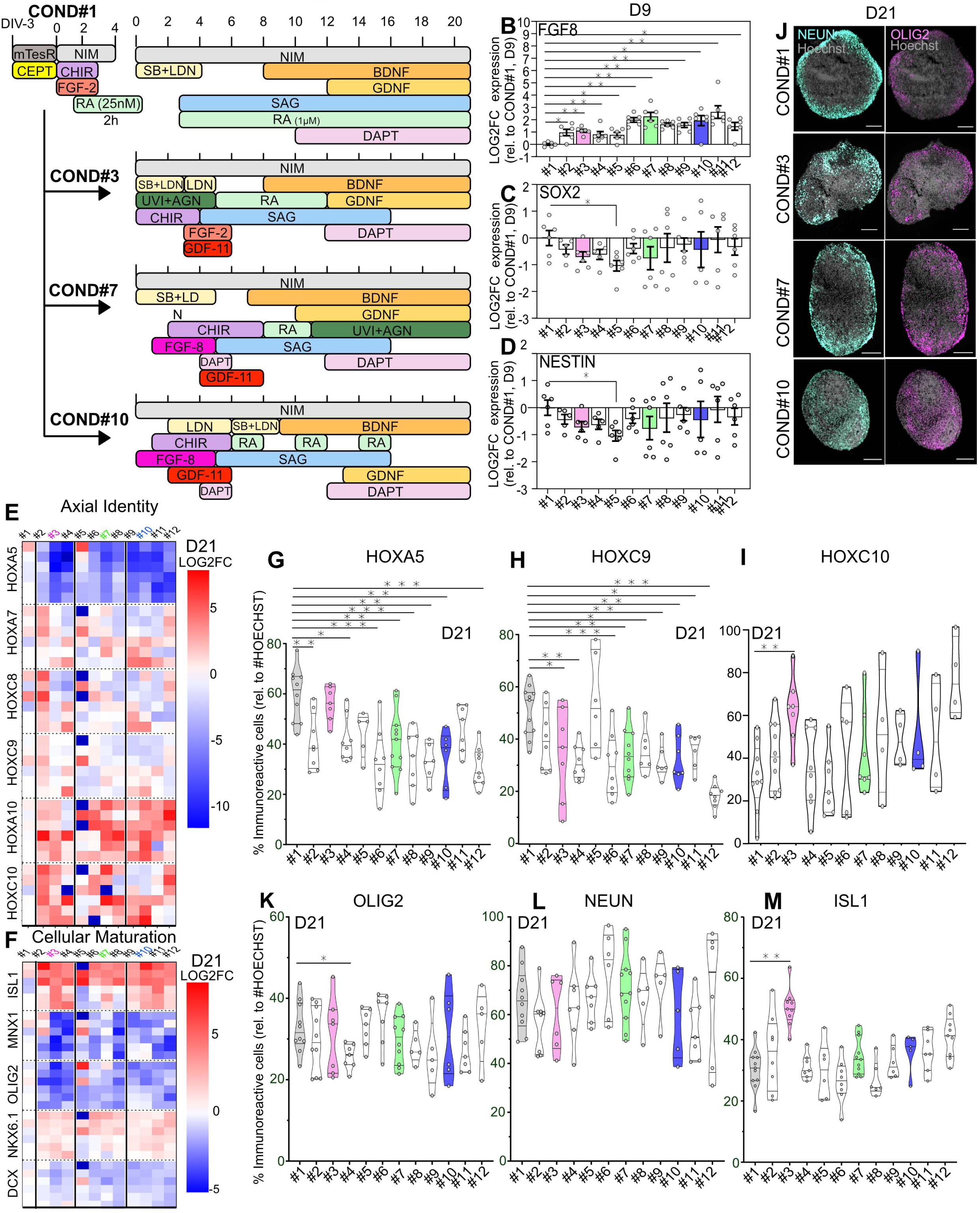
A protocol screen for NMP-derived differentiation of spMNs reveals higher caudalization efficiency in protocols with prolonged NMP state in culture (associated to figure 3) A. Scheme showing the guided differentiation patterning from day 0 to day 21 for standard protocol #1 and the selected protocols per module, all starting from RA-primed NMPs (see Supplemental Table 1). B. qRT-PCR analysis on day 9 spheres generated according to the standard protocol #1 or the new 11 differentiation protocol showing transcript levels the caudal marker *FGF8* (Unparametric Mann-Whitney-U test. N=2 differentiations with n=1-3 technical replicates, each with 5-10 pooled spheres). C. qRT-PCR analysis showing *SOX2* expression on spheres generated as in (B). D. qPCR analysis showing *NESTIN* on spheres generated as in (B). E. Heatmap showing LOG2FC of RC HOX gene expression on day 21 spheres generated according to differentiation protocols #1 to #12. Each row corresponds to a technical replicate generated from 5-10 pooled spheres. N=2 independent differentiation batches, n=6 technical replicates). F. Heatmap showing LOG2FC of neuronal markers gene expression as in (E). G. Quantification of the percentage of HOXA5+ cells from day 21 SCOs generated according to the 12 differentiation protocols. (Unparametric Kruskal-Wallis test with uncorrected Dunn’s test. N=2 differentiations, n=2-5 SCOs/experiment, each dot represents one organoid -average relative cell ratio from 1-3 imaged cryosections-). H. Quantifications of the percentage of HOXC9+ cells from day 21 SCOs generated as in (G). I. Quantifications of the percentage of HOXC10+ cells from day 21 SCOs generated as in (G). J. Representative immunostainings against NEUN (green) and OLIG2 (violet) on day 21 SCOs from the selected key protocols. Scale bar, 200µM K. Quantifications of the percentage of OLIG2+ cells from day 21 SCOs generated as in (G). L. Quantifications of the percentage of NEUN+ cells from day 21 SCOs generated as in (G). M. Quantifications of the percentage of ISL1+ cells from day 21 SCOs generated as in (G). For all graphics, mean±SEM is shown, *p < 0.05, **p < 0.01, ***p < 0.001.

**Supplementary Figure S2.**
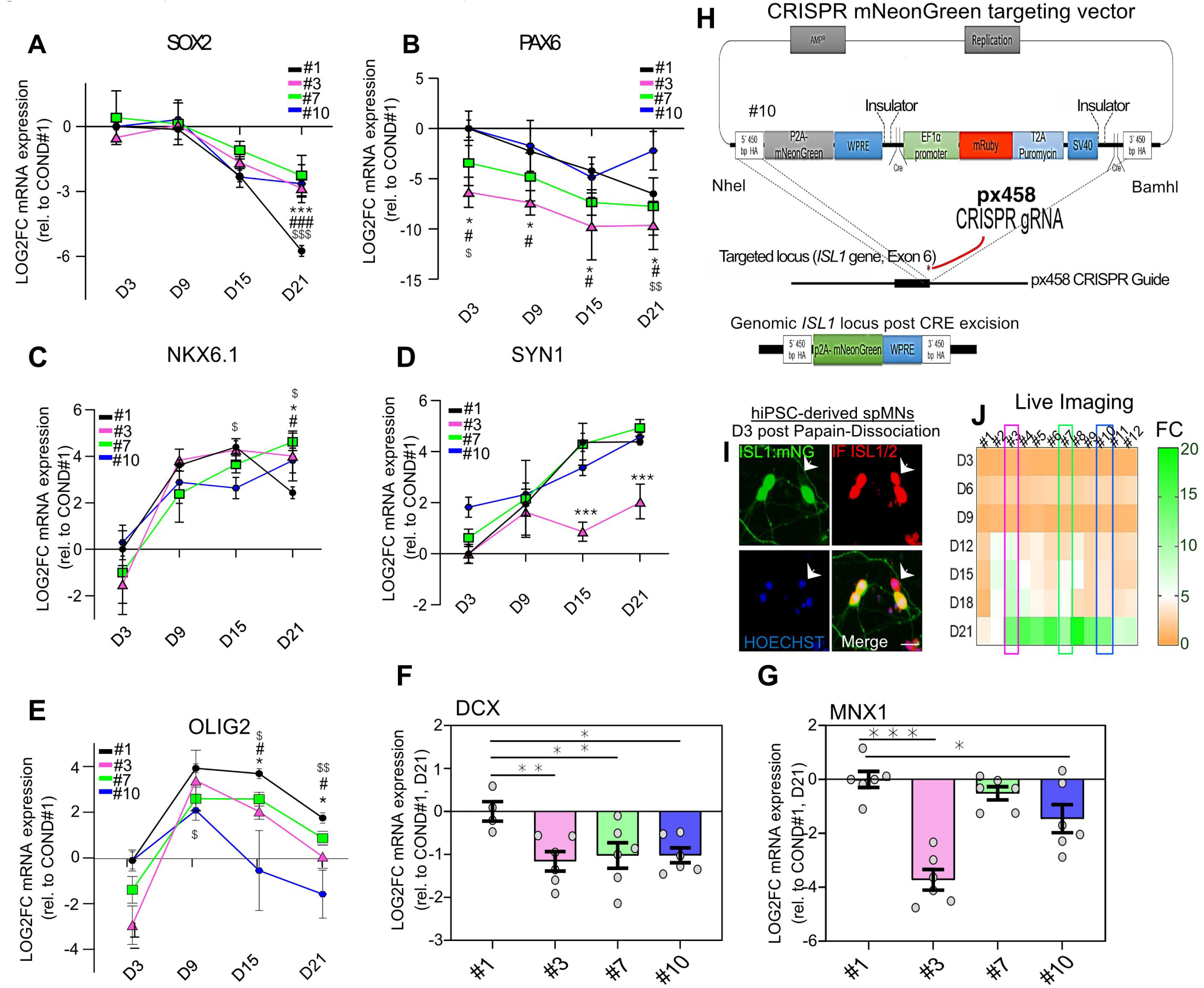
Differences in the longitudinal timeframe of SCO development and spMN generation (associated to figure 3) A. Longitudinal qRT-PCR analysis from day 3, 9, 15 and 21 SCOs generated according to differentiation protocols #1, 3, 7 and 10 showing LOG2FC *SOX2* expression (Unparametric Mann-Whitney U test. N=2 differentiations, n=1-3 technical replicates, each containing 5-10 pooled SCOs. Each dot represents a technical replicate). B. Longitudinal qRT-PCR analysis showing LOG2FC *PAX6* transcript levels from SCOs generated as in (A). C. Longitudinal qRT-PCR analysis showing LOG2FC *NKX6.1* transcript levels from SCOs generated as in (A). D. Longitudinal qRT-PCR analysis showing LOG2FC *SYN1* transcript levels from SCOs generated as in (A). E. Longitudinal qRT-PCR analysis showing LOG2FC *OLIG2* transcript levels from SCOs generated as in (A). F. qRT-PCR analysis showing LOG2FC *DCX* expression from day 21 SCOs generated from the selected protocols. (Unparametric Mann-Whitney U test. N=2 differentiations, n=1-3 technical replicates, each with 5-10 pooled SCOs. Each dot represents a technical replicate). G. qRT-PCR analysis showing LOG2FC *HB9* expression from day 21 SCOs as in (F). H. Scheme showing HR120-p2A-mNeonGreen Targeting vector to C-terminally tag the endogenous *ISL1* locus on the control BJ-siPS hiPSC line using CRISPR-Cas9 and a knock-in approach. I. Representative imaging showing the endogenous ISL1-mNeonG reporter (green) and immunostaining against ISL1/2 (red) on plated spMNs dissociated from BJ-siPS ISL1-mNeonG spinal spheres generated following the differentiation protocol in Fig. 1A. Arrowhead indicates a neuron positive for the antibody staining (against bother ISL1 and ISL2) and negative for the endogenous reporter. J. Heatmap showing fold change of ISL1-mNeonG reporter signal during the SCO development normalized by total SCO area and relative to day 3. SCOs were generated according the 12 different protocol conditions. (N=2 independent differentiations, n=2-8 technical replicates (wells)/experiment, each imaged well contained 5-10 SCOs). For all graphics, mean±SEM is shown, *p < 0.05, **p < 0.01, ***p < 0.001.

**Supplementary Figure S3.**
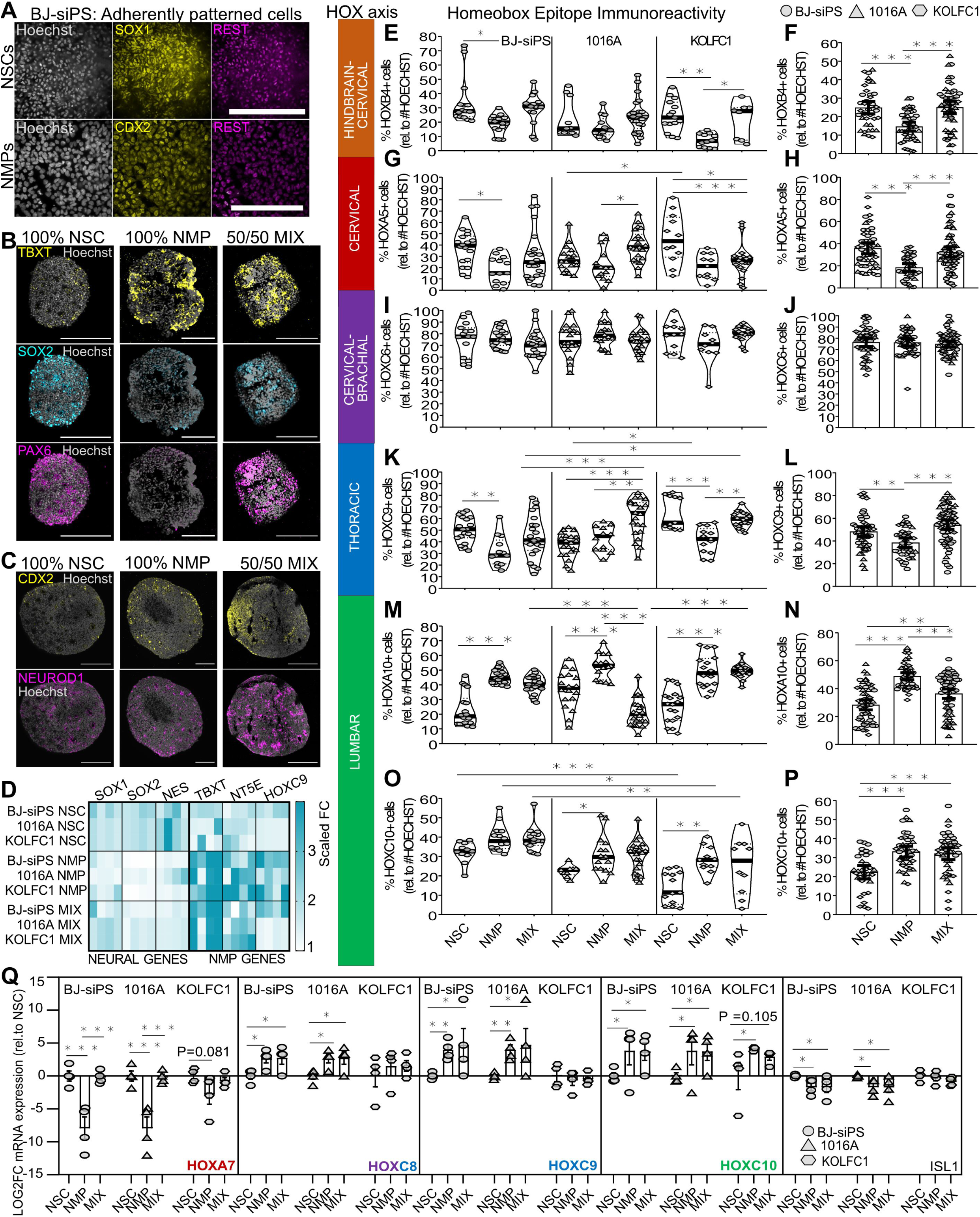
The axial identity of the SCO depends on the stem cell starting population during aggregate assembly (associated to figure 4) A. SOX1 (yellow), CDX2 (yellow) and REST (violet) immunostaining on BJ-siPS-derived adherently patterned NSCs and NMPs. hiPSCs were patterned for 72h in NIM medium+small molecules, according to Fig. 2A. Scale bar, 100µM. B. Immunostained against TBXT (yellow), neural SOX2 (blue) and proneural PAX6 (violet) on day 0 sphere cryosections derived from 100% BJ-siPS NSCs, 100% NMPs or a 50:50 mix of both stem cell types. Scale bar, 200µM. C. Immunostained against caudal marker CDX2 (yellow) and proneural NEUROD1 (violet) on day 0 sphere cryosections derived from 100% BJ-siPS NSCs, 100% NMPs or a 50:50 mix of both stem cell types. Scale bar, 200µM. D. Heatmap showing LOG2FC of qRT-qPCR analysis from day 0 100% NSC, 100% NMP or 50:50%-derived spheres from three healthy hiPSC lines (BJ-siPS, 1016A and KOLFC1). Levels of NSC genes (*SOX1*, *SOX2*, *NES*) and caudal progenitor/NMP markers (*TBXT*, *NT5E*, *HOXC9*) are shown. E. Quantification of the percentage of cells positive for HOXB4 (hindbrain) on day 21 cryosections of SCOs generated from only NSCs, only NMPs or an even mixture of both from the three healthy hiPSC lines. (Two-way ANOVA with Tukey-Kramer test. N = 4-5 differentiations, n = 3- 6 SCOs per experiment. Each dot represents one organoid and is the average of 1-3 imaged cryosections. Violin plots show the median and interquartile range.). F. Quantification of the percentage of HOXB4+ cells as in (E), but values for each stem cell starting population condition from the 3 cell lines are pooled together (One-way ANOVA and FDR). G. Quantification of the percentage of HOXA5+ cells on day 21 SCO cryosections as in (E). H. Quantification of the percentage of HOXA5+ cells on day 21 SCO cryosections as in (F). I. Quantification of the percentage of HOXC6+ cells on day 21 SCO cryosections as in (E). J. Quantification of the percentage of HOXC6+ cells on day 21 SCO cryosections as in (F). K. Quantification of the percentage of HOXC9+ cells on day 21 SCO cryosections as in (E). L. Quantification of the percentage of HOXC9+ cells on day 21 SCO cryosections as in (F). M. Quantification of the percentage of HOXA10+ cells on day 21 SCO cryosections as in (E). N. Quantification of the percentage of HOXA10+ cells on day 21 SCO cryosections as in (F). O. Quantification of the percentage of HOXC10+ cells on day 21 SCO cryosections as in (E. P. Quantification of the percentage of HOXC10+ cells on day 21 SCO cryosections as in (F). Q. qRT-PCR analysis on day 21 SCOs generated from 100% NSCs, 100% NMPs or a 50:50% ratio from three healthy control hiPSC lines showing *HOXA7*, *HOXC8*, *HOXC9*, *HOXC10* and *ISL1* gene expression. (Unparametric Mann-Whitney-U test. N=4 independent differentiations, n=1 technical replicate/experiment, each containing 3 pooled SCOs. Each dot corresponds to one technical replicate). For all graphics, mean±SEM is shown, *p < 0.05, **p < 0.01, ***p < 0.001.

**Supplementary Figure S4.**
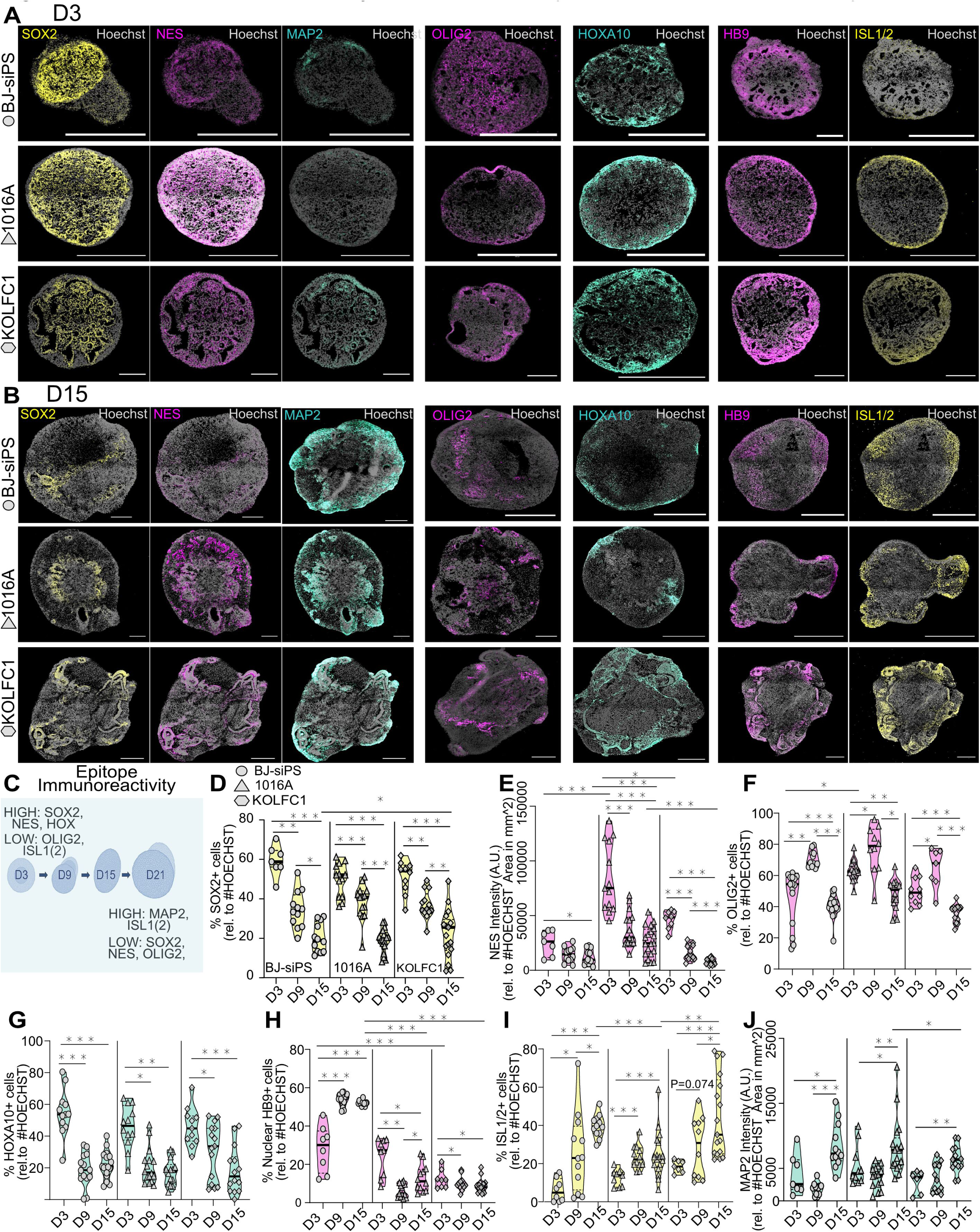
Several hiPSC-lines subjected to the CASCO protocol show similar in vitro development (associated to figure 5) A. Representative immunostaining against SOX2 (yellow), NESTIN (violet), MAP2 (green), OLIG2 (violet), HOXA10 (green), HB9 (violet) and ISL1/2 (yellow) on day 3 CASCOs derived from three healthy hiPSC lines (BJ-siPS, 1016A and KOLFC1). Scale bar, 200µM. B. Representative immunostaining against SOX2 (yellow), NESTIN (violet), MAP2 (green), OLIG2 (violet), HOXA10 (green), HB9 (violet) and ISL1/2 (yellow) on day 15 CASCOs derived from three healthy hiPSC lines (BJ-siPS, 1016A and KOLFC1). Scale bar, 200µM. C. Scheme indicating the markers expected to be detected by immunostaining at each CASCO developmental time point. D. Longitudinal quantification of SOX2+ cells from day 3, 9 and 15 CASCO cryosections. Organoids were derived from the three hiPSC (Two-way ANOVA with Tukey-Kramer test. N=4- 5 differentiations with n=3-6 SCOs per experiment. Each dot represents one organoid, and is the average from 1-3 imaged cryosections. Violin plots show median and interquartile range). E. Longitudinal quantification of NESTIN+ immunostaining relative to the CASCO area stained with HOECHST from samples as in (D). F. Longitudinal quantification of OLIG2+ cells HOECHST from samples as in (D). G. Longitudinal quantification of HOXA10+ cells from samples as in (D). H. Longitudinal quantification of HB9+ cells from samples as in (D). I. Longitudinal quantification of ISL1/2+ cells from samples as in (D). J. Longitudinal quantification of MAP2+ immunostaining relative to the CASCO area stained with HOECHST from samples as in (D). For all graphics, mean±SEM is shown, *p < 0.05, **p < 0.01, ***p < 0.001.

**Supplementary Figure S5.**
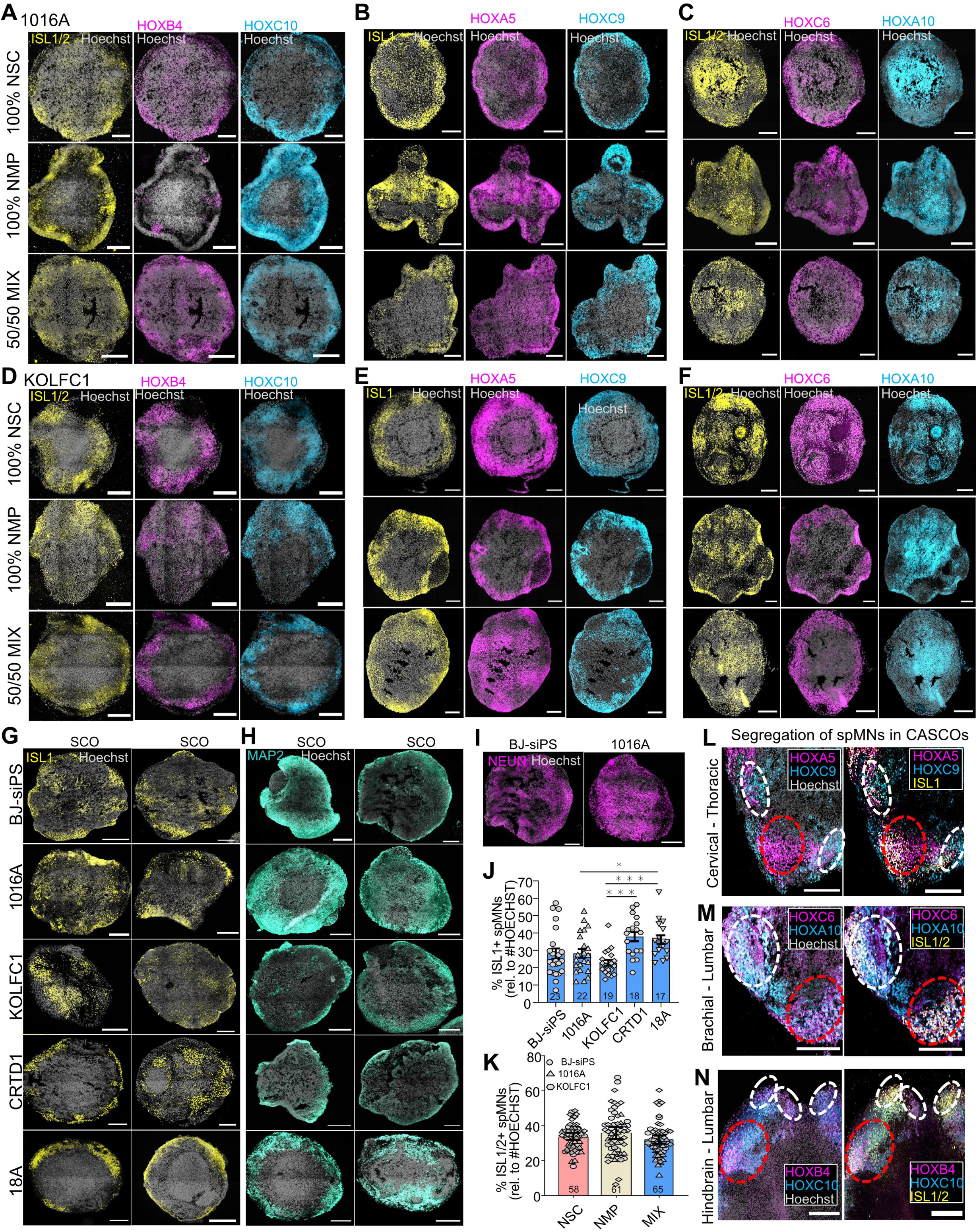
Reproducible generation of CASCOs across cell lines (associated to figure 5) A. Representative immunostaining against ISL1/2 (yellow), hindbrain HOXB4 (violet) and lumbar HOXC10 (blue) on day 21 CASCO cryosections derived from the healthy hiPSC line 1016A under three starting conditions: 100% NSC, 100% NMP, and 50:50% MIX. Scale bar, 200µM. B. Representative immunostaining against ISL1 (yellow), the cervical HOXA5 (violet), the thoracic HOXC9 (blue) on day 21 CASCO cryosections as in (A). C. Representative immunostaining against ISL1/2 (yellow), the brachial HOXC6 (violet), the lumbar HOXA10 (blue) on day 21 CASCO cryosections as in (A). D. Representative immunostaining against ISL1/2 (yellow), hindbrain HOXB4 (violet) and lumbar HOXC10 (blue) on day 21 CASCO cryosections derived from the healthy hiPSC line KOLFC1 under three starting conditions: 100% NSC, 100% NMP, and 50:50% MIX. Scale bar, 200µM. E. Representative immunostaining against ISL1 (yellow), the cervical HOXA5 (violet), the thoracic HOXC9 (blue) on day 21 CASCO cryosections as in (D). F. Representative immunostaining against ISL1/2 (yellow), the brachial HOXC6 (violet), the lumbar HOXA10 (blue) on day 21 CASCO cryosections as in (D). G. Representative immunostaining against ISL1 (yellow) showing two day 21 CASCOs patterned from a 50%:50% mix of hiPSC-derived NSCs or NMPs. Five different healthy hiPSC lines (BJ-siPS, 1016A, KOLFC1, CRTD1, 18A) were used. Scale bar, 200µM. H. Representative immunostaining against the neural filament marker MAP2 (green) on day 21 CASCOs derived from the five healthy hiPSC lines as in (G). Scale bar, 200µM. I. Representative immunostainings against neural maturity marker NEUN (green) from day 21 CASCOs derived from two healthy hiPSC lines patterned as in (G). Scale bar, 200µM. J. Quantification of the percentage of ISL1+ spMNs generated in the CASCOs per hiPSC line, including all experimental batches of the MIX condition. (One-way ANOVA with Uncorrected Dunn’s test. N = 3-5 differentiations with n = 3-6 SCOs per experiment. Each dot represents one organoid and is the average from 1-3 imaged cryosections. Exact number of CASOs indicated in each bar with a number). K. Quantification of the percentage of ISL1/2+ spMNs in the CASCOs generated from the 100% NSC, 100% NMP and 50%:50% conditions and derived from BJ-siPS, 1016A and KOLFC1 hiPSC lines. L. Immunostaining showing zoomed spatial segregation of thoracic cells and spMNs (white dotted line) from cervical cells and spMNs (red dotted line). Showing ISL1 (yellow), HOXA5 (violet), HOXC9 (blue) and Hoechst (grey). Scale bar, 200µM M. Immunostaining showing zoomed spatial segregation of brachial cells and spMNs (red dotted line) from lumbar cells and spMNs (white dotted line). Showing ISL1(2) (yellow), HOXC6 (violet), HOXA10 (blue) and Hoechst (grey). Scale bar, 200µM N. Immunostaining showing zoomed spatial segregation of hindbrain cells and spMNs (white dotted line) from lumbar cells and spMNs (red dotted line). Showing ISL1(2) (yellow), HOXB4 (violet), HOXC10 (blue) and Hoechst (grey). Scale bar, 200µM For all graphics, mean±SEM is shown, *p < 0.05, **p < 0.01, ***p < 0.001.

**Suppl. Table 1.**
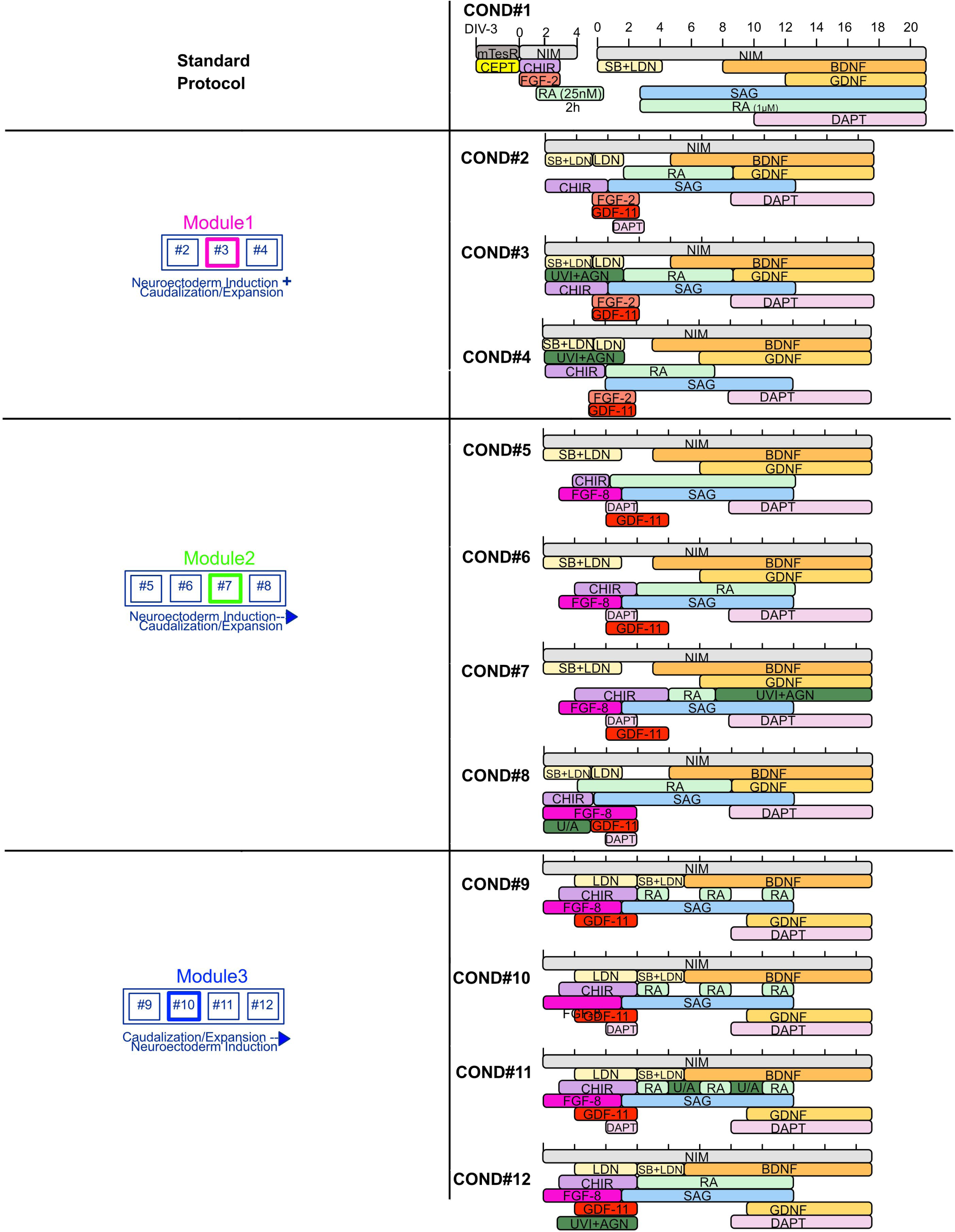
Differentiation protocols of Screen.

